# Mechanisms of ligand recognition and channel opening for P2X2 receptors in lipid nanodiscs

**DOI:** 10.64898/2025.12.04.692396

**Authors:** Surbhi Dhingra, Maia Moog, Kenton J. Swartz

**Affiliations:** Molecular Physiology and Biophysics Section, Porter Neuroscience Research Center, National Institute of Neurological Disorders and Stroke, National Institutes of Health, Bethesda, MD 20892; National Institutes of Health – Karolinska Collaborative Doctoral Program in Neuroscience, Bethesda, MD 20892

## Abstract

Extracellular ATP activates P2X receptor channels (P2XRs) that serve important roles in the immune and nervous systems. Available structures of P2XRs in detergents reveal that ATP binding to the extracellular domain leads to severing of subunit interfaces within transmembrane regions as the pore opens. Here we report cryo-EM structures of the human P2X2R in lipid nanodiscs in an apo closed state, with ATP^4-^, Mg-ATP^2-^ and suramin bound. We find that a unique Arg residue coordinates the γ-PO4 of ATP^4-^ in P2X2R and underlies the requirement of this subtype for ATP^4-^. Channel opening and desensitization occur when ATP^4-^ binds, whereas the channel remains closed when Mg-ATP^2-^ binds. A continuous belt of partially resolved lipids in the outer leaflet stabilizes the closed state, and the presence of lipids prevents the severing of subunit interfaces as the channel opens. These findings establish key mechanistic principles of gating for P2X2R in a membrane-like environment, providing a framework for future mechanistic studies and therapeutic development.

## Introduction

ATP serves as a source of energy throughout biology, but also as a chemical transmitter when released from cells where it activates P2XR channels and P2Y G-protein coupled receptors (*1, 2*). Ionic currents activated by extracellular ATP were first described in sensory (*3*) and spinal cord (*4*) neurons, and subsequently seven subtypes of P2XR subunits were identified and found to be widely expressed where they serve diverse cellular functions (*1, 2*). In sensory neurons, P2XRs are involved in sensing hypoxia in the carotid body (*5*), in taste (*6*), bladder filling (*7*), hearing (*8, 9*) and pain (*1, 10, 11*). In the central nervous system, they are involved in synaptic homeostasis and epilepsy (*1, 12*), in smooth muscle they are involved in vascular contraction (*1, 13*) and in the immune system they participate in inflammation and immunity (*1, 14*).

Structures of P2XRs in detergents (*15–23*) reveal that these trimeric cation channels contain a large extracellular domain (ECD) with binding sites for ATP at subunit interfaces and two TM helices in each subunit that form the ion conduction pore. The TM2 helix lines the pore and contains the activation gate that prevents ion permeation when closed (*15–27*). These structures have provided an indispensable framework for understanding P2XRs, yet many fundamental questions remain to be answered. For example, one of the enigmatic features of the available P2XR structures in detergents where open states have been resolved is the severing of subunit interfaces, creating large intramembrane crevices (*15–19*). These intersubunit crevices are large enough that lipids exchange between the membrane and the aqueous ion permeation pathway in Molecular Dynamics (MD) simulations to clog the pore and prevent ion permeation (*24*). Lipids might organize around these crevices to prevent them from entering the pore or these crevices might be a non-native feature resulting from the use of detergents. Thus, an important and outstanding question is whether the presence of a lipid membrane plays a role in shaping the structure and dynamics of the pore during the gating cycle of P2XRs. The key structural elements coupling ATP binding to channel opening also remain to be identified.

The P2X2R subtype has been extensively studied and many important physiological functions have been described using electrophysiological approaches (*1, 2, 28*). P2X2R requires ATP^4-^ for activation (*19, 29, 30*), whereas some other subtypes can be activated by the physiologically abundant divalent-cation bound forms of ATP (*19, 30*). For example, P2X3Rs contain an acidic chamber that stabilizes a Mg^2+^ ion bound to ATP that is required for activation by Mg-ATP^2-^ (*19*). In contrast, for P2X2Rs the structural basis for requiring ATP^4-^ and how the channel opens upon binding of the nucleotide remain to be elucidated.

In the present study we report structures of the hP2X2R, using lipid nanodiscs to understand the role of lipids in shaping the structure of the pore. We succeeded in capturing key conformations in the gating cycle of P2X2Rs, revealing that membrane lipids stabilize the subunit interfaces as the channel opens. We also show how hP2X2 distinguishes between ATP^4-^ and Mg-ATP^2-^, and how suramin functions as a competitive antagonist.

## Results

### Structure of apo hP2X2 in lipid nanodiscs

hP2X2R has two splice variants that exhibit similar functional properties (*31*), with the longer variant (hP2X2-L) containing an additional 67 residues within the C-terminus missing from the shorter variant (hP2X2-S) (**Fig. S1**). We began by expressing hP2X2-S in mammalian cells and reconstituting the purified protein into lipid nanodiscs with Na^+^ as the only cation to capture the apo state. We used cryogenic electron microscopy (cryo-EM) to evaluate the structure of hP2X2-S, however, refinement using C3 symmetry revealed extra density within the ATP binding site (*15–20*), suggesting that cellular ATP co-purified with the protein (*17*). To obtain an apo structure of the P2X2 receptor, we removed ATP during cell lysis by incubating the protein with apyrase and alkaline phosphatase and determined structures of both apo hP2X2-S (3.1 Å) and apo hP2X2-L (2.7 Å) reconstituted into nanodiscs (**Fig. S3; Table S1, S2**). We focus on the description of apo hP2X2-L, but we have not identified any differences between the two variants.

The structure of apo hP2X2-L has an architecture (**Fig. 1a,b**) similar to other subtypes (*15–23*), with each subunit resembling a dolphin (**Fig. 1c**). The homotrimeric receptor contains a large ECD and a relatively small TM domain (TMD), and contains two glycosylation sites (N194 and N310) seen in other subtypes and one unique site (N133) (**Fig. 1a-c,j**). The quality of our cryo-EM map was sufficient to resolve side chain densities for most residues (**Fig. S3**). Within the ECD, hP2X2-L differs from other subtypes within the head domain involved in ATP binding. Previous structures contain a loop in the head domain whereas hP2X2-L contains an additional β-strand (β-9) (**Fig. 1j,k; Fig. S2**). The unique head domain of hP2X2-L also contains a well-resolved Arg residue (R130) directed into the ATP binding pocket (**Fig. 1j**) that is not present in other subtypes (*17, 18, 20, 21, 23*) (**Fig. S2**). External lateral fenestration are present between the ECD and the TMD (**Fig. 1a,b**), consistent with functional studies on rat P2X2-L (rP2X2-L) suggesting that ions permeate through these lateral fenestrations (*25, 32*).

**Fig. 1.**
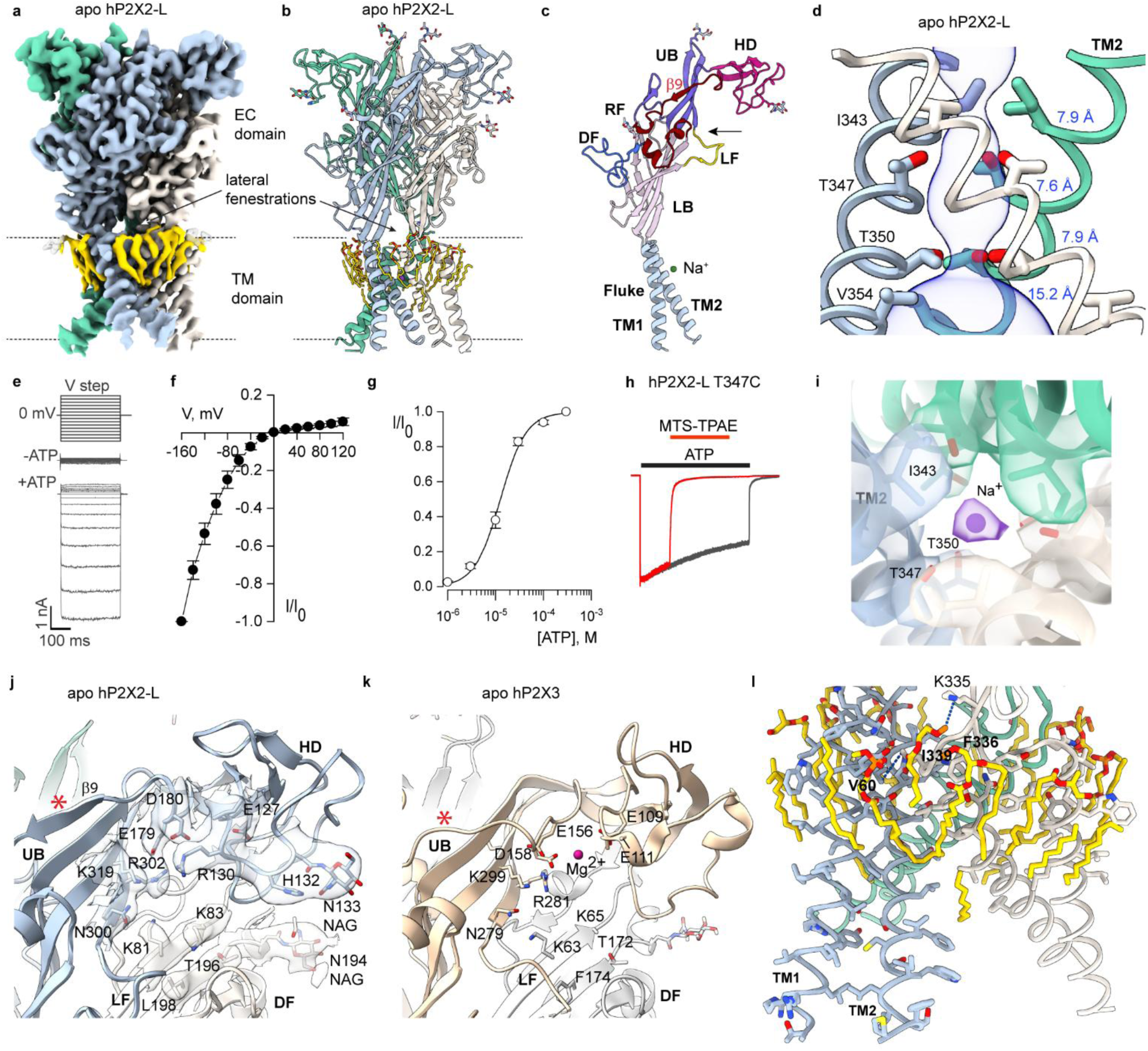
Structure of the apo hP2X2R. **a**) Cryo-EM map of apo hP2X2-L (CryoSPARC sharpened) with different subunits colored in steel blue, aquamarine and linen, and lipids in gold. **b**) Structural model of apo hP2X2-L. **c**) Single subunit of apo hP2X2-L illustrating the dolphin-like shape with regions corresponding to body parts color coded and labeled as originally proposed based on the structure of P2X4 (*15*). The unique β9 strand in hP2X2-L has been labelled in red. Abbreviations for body parts are Upper Body (UB), Head Domain (HD), Left Flipper (LF), Dorsal Fin (DF) and Lower Body (LB). **d**) Structure of the activation gate region made up of the three TM2 helices in the pore of apo hP2X2-L shown as a side view along with MOLE (*73*) representation. Residues in the regions of the pore that function as gates are shown as stick representation with distances shown in blue between adjacent Cα atoms for I343, T347, T350 and V354. **e**) Currents elicited by applying voltage steps from −160 to +120 mV in 20-mV increments from a holding potential of 0 mV for hP2X2-L in the absence and presence of 300 µM ATP. **f**) Normalized plots of current-voltage relationships for ATP activated currents obtained by subtracting control currents recorded in the absence of ATP from the currents recorded in the presence of ATP. The data points are mean steady-state current ± SEM (n=3 in 2 independent experiments), normalized to ATP activated-current at −160 mV. **g**) Normalized concentration-response relations for ATP activation of hP2X2-L in a physiological solution containing divalent cations (see Methods) (n=9 in 3 independent experiments). The solid lines represent fits of the Hill equation to the data resulting in an EC_50_ = 16.8 ± 1.8 μM and nH = 1.3 ± 0.2. **h**) MTS-TPAE rapidly reacts with T347C in hP2X2-L in the presence of 300 µM ATP and nearly completely inhibits the current. The fraction of current remaining after reaction with MTS-TPAE was 0.022 ± 0.008 for T347C in hP2X2-L (n=4 in 2 independent experiments), very similar to the value of 0.023 ± 0.015 (n=4 in 2 independent experiments) for reaction of MTS-TPAE with the equivalent residue (T336C) in rP2X2-L. **i**) C1 cryo-EM map (CryoSPARC sharpened, contour ∼0.13) and model generated from the C3 map for the gate region of apo hP2X2-L showing density corresponding to a Na^+^ ion bound within the gate and interacting with the Oγ of T347. Evaluation of apo hP2X2-L using CheckMyMetal resulted in an acceptable occupancy of 1. **j**) Structure of apo hP2X2-L around the agonist recognition pocket. Cryo-EM density (CryoSPARC sharpened, contour ∼0.04) is shown for key side chains near to where ATP binds (see Fig. 2). Red asterisk indicates the presence of the unique β9 strand. **k**) Structure of apo hP2X3 around the agonist recognition pocket (5svj) (*17*). **l**) TM region of hP2X2-L showing key residues that interact with lipids in the outer half of the lipid bilayer and those involved in packing between TM1 and TM2.

Like other P2XRs, each subunit contains two TM helices, with TM2 lining the pore at the central axis with two residues (I343 and T350) just external to the middle of the membrane occluding the ion permeation pathway (**Fig. 1d**), suggesting that it represents a closed state. The location of an activation gate between I343 and T350 is supported by cysteine accessibility studies in rP2X2-L showing that modification by thiol reactive compounds occurs equally rapidly at the equivalent of I343 in closed and open states, becomes progressively slower in the closed state at positions equivalent to T347 and T350, and that larger thiol reactive compounds abolish ion permeation when reacted at the equivalent of T347C (*26, 27*). To confirm that the functional properties of hP2X2-L are similar to rP2X2-L, we expressed the channel in HEK cells and used whole-cell patch clamp recordings to measure macroscopic ionic currents. We observed robust inwardly-rectifying current (I)-voltage (V) relationships when applying external ATP, similar to rP2X2-L (*33–35*) (**Fig. 1e,f**). The apparent affinity for ATP in the presence of physiological concentrations of divalent cations was also similar to rP2X2-L (**Fig. 1g**) (*27*) and hP2X2-L can be completely inhibited when a large thiol-reactive compound reacts with the T347C mutant within the gate (**Fig. 1h**), suggesting that key functional properties of hP2X2-L are similar to rP2X2-L (*26, 27*).

The TMD of apo hP2X2-L in lipid nanodiscs has similar overall architecture to that found in other subtypes, however the TM1 and TM2 helices are somewhat more tightly packed (**Fig. 1l**). We attempted to determine the structure of apo hP2X2-L in the detergent n-dodecyl-β-d-maltopyranoside (DDM) to compare with the structure in nanodiscs but found that ATP co-purified with the protein and could not be completely removed with apyrase and alkaline phosphatase (**Fig. S3, S4c; Table S1, S2**). The two structures we solved in detergent (without and with apyrase/alkaline phosphatase) are distinct from the apo structure in nanodiscs (**Fig. S4d,e**), likely due to residual ATP. A recent structure of hP2X2 in DDM with cholesteryl hemisuccinate did not contain residual ATP (*22*) and is remarkably similar to our apo structure in nanodiscs (backbone RMSD of 0.7 Å), suggesting that the presence of lipids does not appreciably alter the structure of apo hP2X2. The structure of apo hP2X2-L in nanodiscs also contains a continuous belt of partially resolved lipids within the outer leaflet hugging the TMD, some with sufficient resolution to see both acyl chains (**Fig. 1a,l**). The headgroups of these lipids are not resolved, so we modeled them as POPC; in one instance the headgroup is positioned near a basic residue (K335) and another near to F336 (**Fig. 1l**). The apo structure of hP2X2-L also reveals interactions between V60 in TM1 in one subunit and I339 in TM2 in the adjacent subunit (**Fig. 1l**). The V60L mutant causes hearing loss in humans (*8*) and its interaction with I339 is important for stabilizing the closed state (*36–39*) and in our structure the V60/I339 interaction is positioned close to the lipids within the outer leaflet (**Fig. 1l**). These observations suggest that the lipid membrane likely plays an important role in stabilizing the structure of P2X2 receptors in the closed state.

The structure of the activation gate in hP2X2-L is interesting in that I343 and T350 constrict the internal and external ends of the gate (0.8 Å and 0.9 Å radii), respectively, leaving a conspicuous cavity (2.2 Å radius) between these residues (**Fig. 1d**). We observed density within this cavity that we attributed to a Na^+^ ion (**Fig. 1i**) as this is the only cation present in our solutions. In apo zebra fish P2X4 (*15*), the residues equivalent to I343, T347 and T350 form a continuous hydrophobic plug. However, cavities within the gate are also seen in apo structures of hP2X1 (*20*) and hP2X3 (*17*), and in the latter a Na^+^ ion was modeled into the cavity. The gate region in rP2X7 is occupied by different residues, but a Na^+^ ion was modeled in this region in the closed state (*40*). We speculate that Na^+^ occupancy of this cavity within the gate of P2XRs helps to stabilize the channel in a closed state, with side chain Oγ atoms of T347 interacting with the ion for hP2X2. Extensive single channel recordings with hP2X7 have shown that substitution of Na^+^ with larger cations progressively increases the open probability of the channel by prolonging the mean open time (*41*), consistent with binding of Na^+^ within the activation gate stabilizing the closed state.

### Mechanisms of ligand recognition

One unique feature of P2X2Rs is that they can only be activated by ATP^4-^ (*30*). ATP bound by the divalent cations Mg^2+^ or Ca^2+^ in physiological solutions can bind to the agonist binding site but cannot open the channel with detectable probability (*30, 42*). To understand the unique mechanism of agonist recognition in P2X2Rs, we determined a series of structures of hP2X2-L or hP2X2-S in nanodiscs with ATP^4-^, Mg-ATP^2-^, Mg^2+^ alone or with the antagonist suramin bound. For several of these structures we also included PIP2 as this lipid slows desensitization (*43*). The inward-rectification seen in I-V relations for hP2X2-L receptors (**Fig. 1e,f**) results from strong rectification of the single channel conductance, but also from a lower open probability at positive voltages (*33, 34*). Since our structures were determined at 0 mV, we used focused classification around the TMD to distinguish particles in different conformations (**Fig. S5**).

By briefly (2.5 s) applying ATP^4-^ (100 µM) with EDTA (5 mM) to chelate all divalent ions, we determined a series of cryo-EM structures of hP2X2-L with ATP^4-^. After focused classification and local refinement imposing C3 symmetry, we observed four classes resolved with comparable resolutions of ∼2.6 Å (**Fig. S5; Table S3**). The density for the TMD was sufficiently well-resolved to model most side chains, in particular within the external leaflet of the membrane. Diffuse density likely corresponding to the cytoplasmic cap at the intracellular side of the TMD like that in structures of P2X3, P2X4 and P2X7 (*17, 18, 23*) could be observed in classes 1, 2 and 4 (**Fig. 2a,b; Fig. S5**), suggesting that the cap is likely present (*44*) even though it is not well-resolved in any of our structures. Similar to apo hP2X2-L, the cryo-EM maps for the ECDs in all four classes with ATP^4-^ bound had local resolutions between ∼2.2 - 2.8 Å, enabling us to see side chain densities for most residues (**Fig. S5**).

**Fig. 2.**
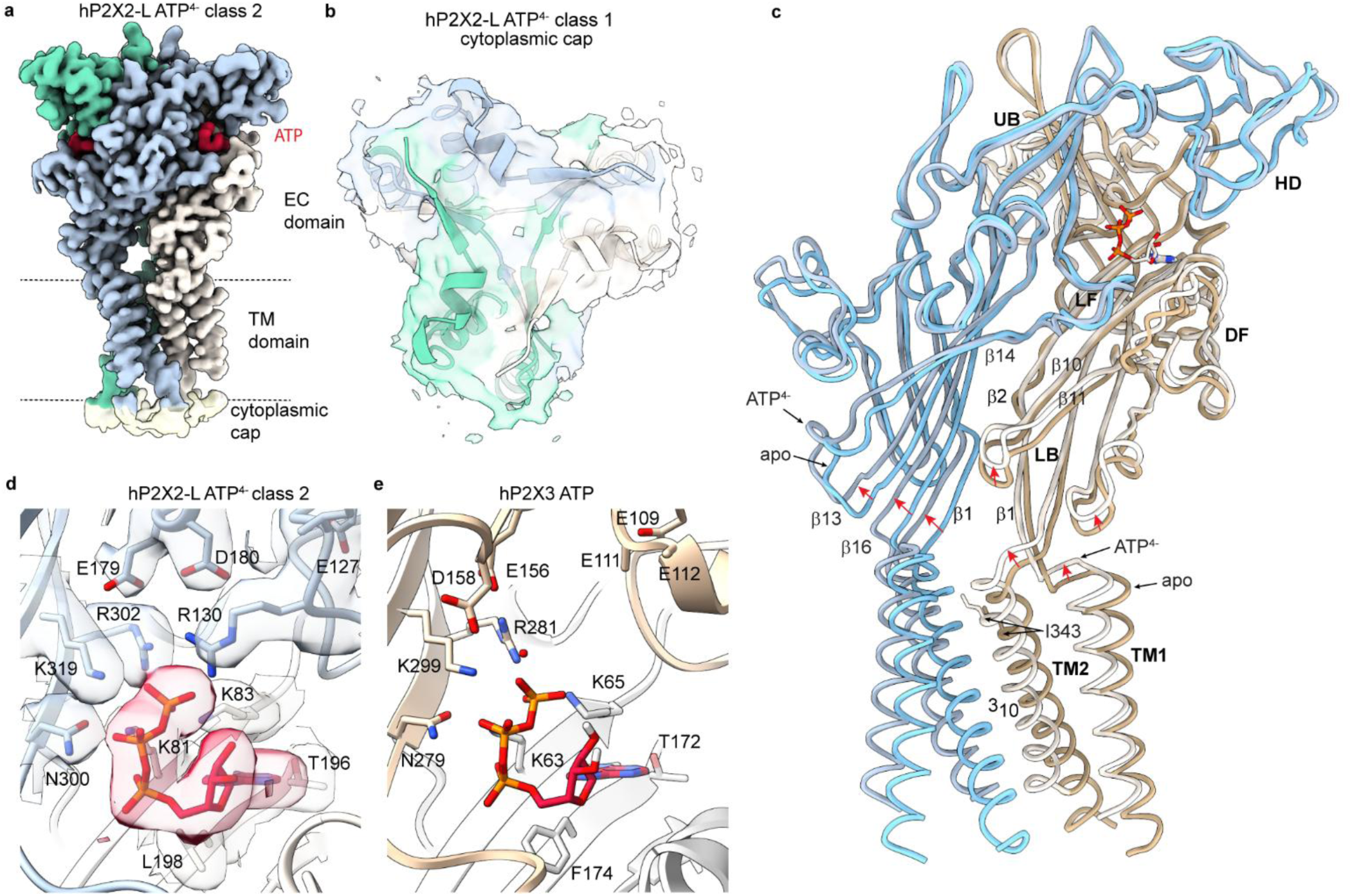
Structure of hP2X2R with ATP^4-^ bound. **a**) Cryo-EM map (DeepEMhancer sharpened) for class 2 of hP2X2-L with ATP^4-^. Density for ATP is shown in red. **b**) Cryo-EM density (unsharpened, contour ∼0.04) for the cytoplasmic cap region of class 1 of hP2X2-L with ATP^4-^ bound from an intracellular perspective. An AlphaFold3 model of hP2X2 is shown fitted into the cryo-EM map. **c**) Superimposed models for apo hP2X2-L and with ATP^4-^ bound (class 2). **d**) Structure of hP2X2-L with ATP^4-^ bound within the agonist recognition pocket. Cryo-EM density (DeepEMhancer sharpened, contour ∼0.02) is shown for key side chains near to where ATP binds. **e**) Structure of hP2X3 with ATP bound within the agonist recognition pocket (5svk) (*17*). Red sphere denotes a water molecule.

To evaluate the ATP site in hP2X2, we focused on class 2, though all classes were similar in the ECD (**Fig. S6a**). ATP^4-^ binds to a pocket formed at the subunit interface, adopting a pose similar to other subtypes (*15–20*) where the nucleotide interacts with residues in the head domain (HD), upper body (UB) and lower body (LB) (**Fig. 2a,c,d**). Comparison of the ATP^4-^ bound and apo structures reveal that agonist binding produces modest conformational changes around ATP (**Fig. 2c; Movie S1**), including an upward movement of β2 and β10 strands in the LB that contain residues interacting with ATP (K81 and K83 in β2 and T196 and L198 in β10) (**Fig. 2c,d; Movie S1**). The dorsal fin (DF) connected to β11 in the LB also moves upward, consistent with a metal bridge between the HD and DF that stabilizes the open state in rP2X2 (*45*), and we see a subtle outward flexing of the L-flipper (LF) (**Fig. 2c**). These movements are similar to those observed in X-ray structures of hP2X3Rs of considerably lower local resolution where many side chains were not modeled (*17*). Binding of ATP^4-^ to hP2X2-L produces more substantial conformational changes in the β-strands of the LB that connect the ECD with the TMD (**Fig. 2c; Movie S1**). The TM1 and TM2 helices move upward by one helical turn, rotate in the counterclockwise direction when viewed from the extracellular side and move laterally to open the channel (see below).

One interesting feature of hP2X2-L bound by ATP^4-^ is the presence of R130 within the HD interacting with the γ-PO_4_ of ATP^4-^ (**Fig. 2d**), which contributes to reducing the size of the binding pocket compared to other subtypes (**Fig. 2d,e; Fig. S6b-d**) and suggesting that the structure of hP2X2 will be useful for developing therapeutics that selectively target this subtype. One robust pharmacological difference between P2X2 and other subtypes like P2X3 is that the non-hydrolyzable and less flexible analogue α,β-Methylene-ATP is an efficacious agonist for P2X3 but not P2X2 (*46, 47*). We tested whether R130 might contribute to this robust difference by mutating it to Ala but observed that α,β-Methylene-ATP could only weakly activate either WT hP2X2 or the R130A mutant (**Fig. S6e-f**), suggesting that other differences are likely involved in determining sensitivity to this agonist.

To investigate the functional impact of R130 in hP2X2, we measured the concentration-dependence for activation by ATP for WT and both R130A and R130E mutants in the presence of divalent cations at concentrations (2 mM Ca^2+^, 0.5 mM Mg^2+^) similar to physiological solutions. The R130A mutant exhibited a modest decrease in apparent affinity for ATP under these conditions while the R130E mutant was indistinguishable from WT (**Fig. 3a**). Given the interaction between R130 and the γ-PO_4_ of ATP^4-^ we see in our structures, we expected that both mutants would weaken ATP interaction with the receptor. We reasoned that the presence of divalent ions in this initial experiment might enable the R130E mutant to interact with the γ-PO_4_ of ATP via a divalent ion, thus effectively rescuing the loss of an interaction between R130 and the γ-PO_4_ of ATP. To test this hypothesis, we measured the concentration-dependence for activation of WT and the two R130 mutants in the presence of 10 mM EDTA, a procedure that is challenging due to the absence of divalent ions that stabilize the seal between the glass pipette and the cell membrane (*48*). Consistent with previous results (*30*), the removal of divalent ions shifted the concentration-dependence for activation by ATP by about an order of magnitude to lower ATP concentrations (**Fig. 3b**), reflecting the fact that only ATP^4-^ can appreciably activate P2X2 (*30*). Under these divalent-free conditions, both R130A and R130E shifted the concentration-dependence for activation by ATP to higher concentrations to similar extents (**Fig. 3b**), consistent with their weakening the binding of ATP. We conclude that the interaction between R130 with the γ-PO_4_ of ATP we observed in our structures stabilizes ATP on the receptor.

**Fig. 3.**
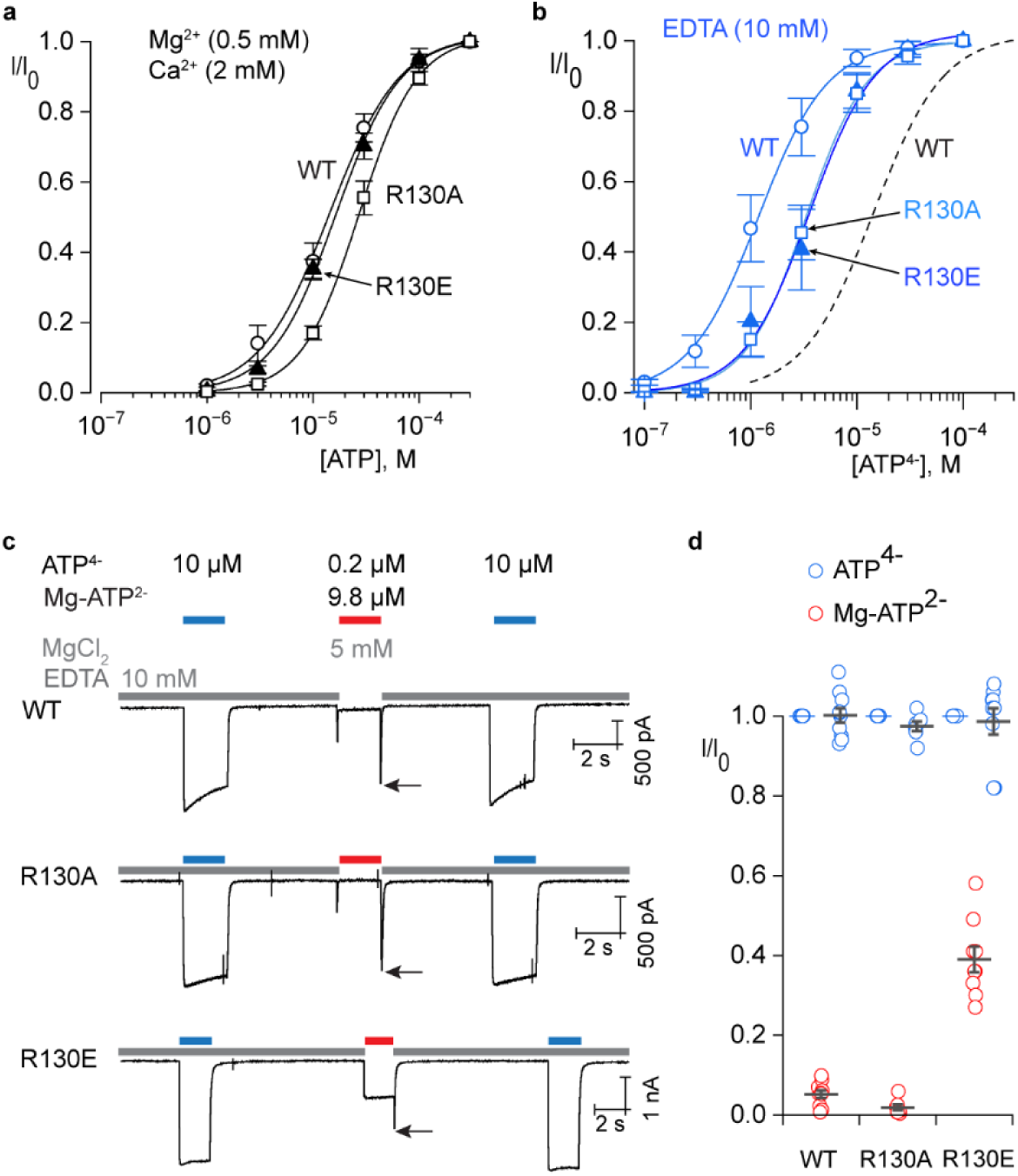
Influence of R130 on agonist recognition in hP2X2. **a**) Normalized concentration-response relations for ATP activation of hP2X2-L WT (n=9 in 3 independent experiments), R130A (n=10 in 3 independent experiments) and R130E (n=10 in 3 independent experiments) in a physiological solution containing the indicated concentrations of divalent cations. Smooth lines represent fits of the Hill equation to the data resulting in an EC_50_ = 16.8 ± 1.8 μM and nH = 1.3 ± 0.2 for WT, EC_50_ = 28.8 ± 3 μM (p=0.0038) and nH = 1.6 ± 0.1 for R130A and EC_50_ = 18.2 ± 2.5 μM (p=0.6392) and nH = 1.4 ± 0.2 for R130E. **b**) Normalized concentration-response relations for ATP activation of hP2X2-L WT (n=6 in 2 independent experiments), R130A (n=5 in 2 independent experiments) and R130E (n=5 in 2 independent experiments) in a solution containing EDTA. Smooth lines represent fits of the Hill equation to the data resulting in an EC_50_ = 1.4 ± 0.4 μM and nH = 1.3 ± 0.3 for WT, EC_50_ = 3.7 ± 0.9 μM (p=0.039) and nH = 1.6 ± 0.1 for R130A and EC_50_ = 4.0 ± 0.9 μM (p=0.021) and nH = 1.8 ± 0.1 for R130E. Dashed black line is the fit of the Hill equation to WT data obtained with divalent ions from panel a for comparison. **c**) Currents elicited at a holding voltage of −60 mV for cells expressing WT hP2X2-L, R130A and R130E when applying external solutions containing 10 µM ATP^4-^ (in EDTA) or 10 µM ATP plus 5 mM MgCl_2_ (containing 0.2 µM ATP^4-^ and 9.8 µM Mg-ATP^2-^). Black arrows identify partially resolved resurgent currents (*30*) that activate when switching from the solution containing Mg-ATP^2-^ back to the control EDTA solution. These resurgent currents result from Mg^2+^ ions unbinding more rapidly than ATP^4-^, enabling the more efficacious form of the bound agonist to transiently activate the channel before unbinding (*30*). Smaller transients observed when first applying Mg-ATP^2-^ are irregular artifacts seen with rapid solution exchange. **f**) Plots of normalized currents activated by ATP^4-^ before and after those activated by Mg-ATP^2-^ for WT and two R130 mutants. Individual cells are shown as open symbols with mean and S.E.M. shown as bars with n=10 for WT, n=7 for R130A and n=9 for R130E, each obtained in three independent experiments.

If divalent ions can mediate an interaction between R130E and the γ-PO_4_ of ATP, we wondered whether this mutant might enable P2X2 to be activated by divalent bound ATP, similar to the influence of acidic residues near to the γ-PO_4_ of ATP in hP2X3 that are critical for activation by divalent ion bound ATP (*19*) (**Fig. 2e**; **Fig. 4e**). To explore this possibility we activated hP2X2 with a near saturating concentration of ATP^4-^ (10 µM) in the presence of 10 mM EDTA and then applied 10 µM ATP together with 5 mM Mg^2+^, a solution that contains 9.8 µM Mg-ATP^2-^ and only 0.2 µM ATP^4-^, a concentration of ATP^4-^ that produces less than 10% activation (*30*) (**Fig. 3c,d**). Even though Mg-ATP^2-^ cannot effectively activate P2X2Rs, we know that divalent ion bound ATP can bind to the receptor because a resurgent current can be resolved when rapidly exchanging the Mg-ATP^2-^ solution for one containing EDTA (*30*). This resurgent current results from Mg^2+^ unbinding more rapidly than ATP, transiently leaving ATP^4-^ on the receptor to open the channel (*30*) (**Fig. 3c**). Mg-ATP^2-^ binding to P2X2Rs without opening the channel has also been demonstrated using a fluorescent ATP analogue (*42*). For both WT and R130A, activation by Mg-ATP^2-^ was negligible and resurgent currents were observed when switching from Mg-ATP^2-^ to EDTA solutions (**Fig. 3c**; black arrows). In contrast, for the R130E mutant, Mg-ATP^2-^activated the channel robustly (**Fig. 3c,d**). A detectable resurgent current was also observed for R130E (**Fig. 3c**; black arrow), indicating that the open probability for Mg-ATP^2-^ remains lower than that for ATP^4-^. We conclude that the interaction between R130 in hP2X2 and the γ-PO_4_ of ATP underlies the requirement of this subtype for ATP^4-^, and that substitution with an acidic residue enables activation by Mg-ATP^2-^.

**Fig. 4.**
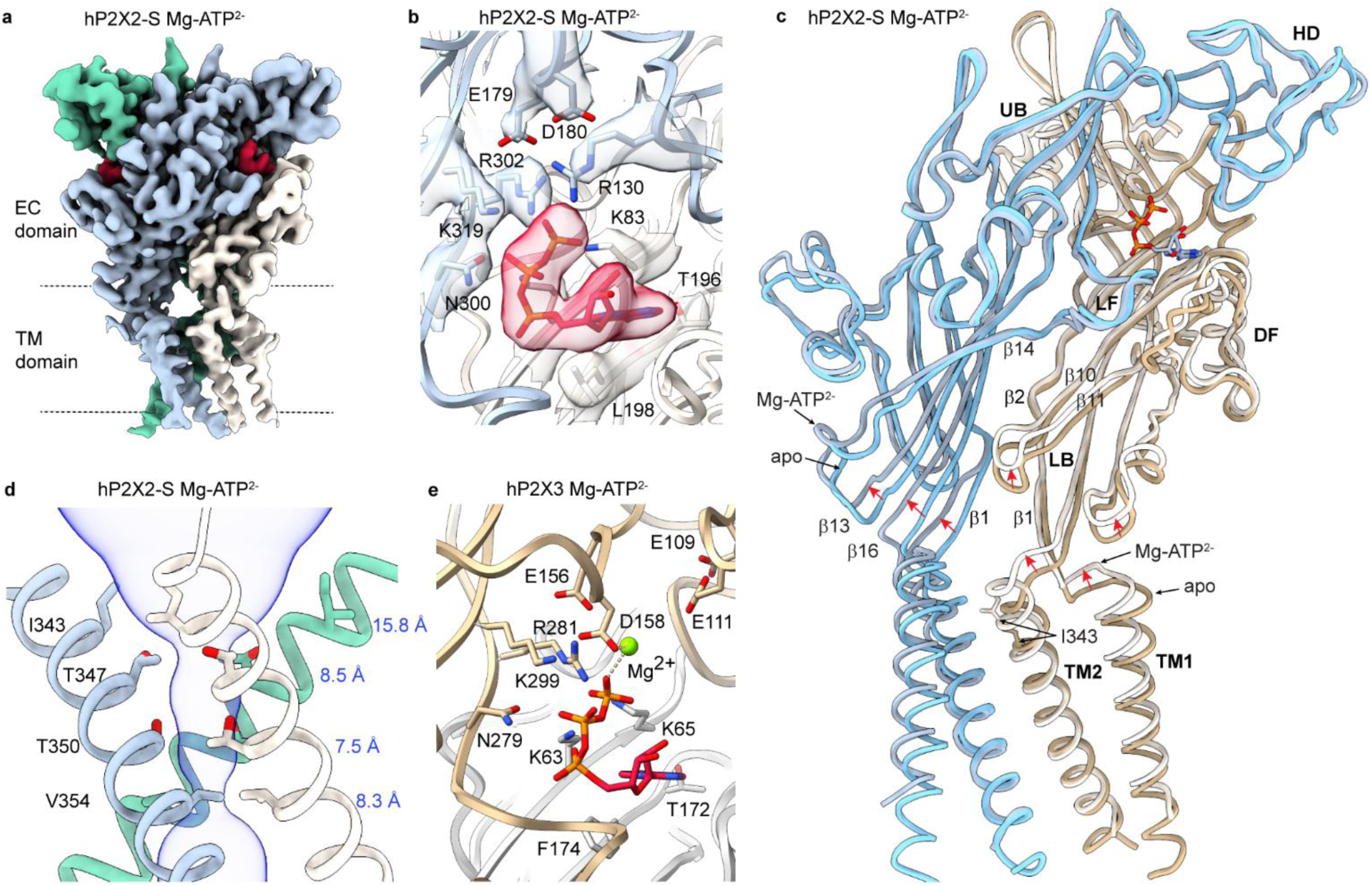
Structure of the hP2X2R with Mg-ATP^2-^ bound. **a**) Cryo-EM map (CryoSPARC sharpened) for hP2X2-S with Mg-ATP^2-^ bound. Density for ATP is shown in red. **b**) Structural model for hP2X2-L with ATP^4-^ bound within the agonist recognition pocket with cryo-EM density (CryoSPARC sharpened, contour ∼0.05) shown for key side chains around ATP. **c**) Superimposed models for apo hP2X2-S and with Mg-ATP^2-^ bound. **d**) Structure of the activation gate region for Mg-ATP^2-^ hP2X2-S shown as a side view along with MOLE (*73*) representation. Residues in the regions of the pore that function as gates are shown as stick representation with distances shown in blue between adjacent Cα atoms for I343, T347, T350 and V354. **e**) Structural model of hP2X3 with Mg-ATP^2-^ bound (6ah4).

To explore why the binding of divalent-bound ATP fails to open hP2X2Rs, we determined a cryo-EM structure of hP2X2-S with Mg-ATP^2-^. After 3D classification using a focused mask for the ATP binding pocket, local refinement imposing C3 symmetry yielded a class with ATP bound at a resolution of 2.9 Å (**Fig. 4a; Fig. S7; Table S1**). Although we could not resolve unambiguous density for Mg^2+^ within the agonist binding pocket (**Fig. 4b**), we observed a substantial conformational change within the ECD, with the largest movements occurring towards the lower end of the β-strand network near the membrane (**Fig. 4c; Fig. S6g,h**). Both TM1 and TM2 translate in the extracellular direction, TM2 rotates counterclockwise when viewed from the extracellular side as in the structure with ATP^4-^ (but to a lesser extent), while the TM1 helix rotates in a clockwise direction, opposite to what we observed for ATP^4-^ (**Fig. 2c**; **Fig. 4c; Fig. S8h;** see discussion). The outermost residue in the gate (I343) rotates outwards and no longer narrows the pore, but the gate remains narrow at T347, T350 and V354 (1.7 Å, 1.8 Å and 0.9 Å radii, respectively) (**Fig. 4d**). These important structural differences between hP2X2 with ATP^4-^ and Mg-ATP^2-^, together with the functional evidence described above (*30*) that Mg-ATP^2-^ can bind to hP2X2Rs but cannot open the channel (**Fig. 3c,d**), support the conclusion that Mg^2+^ ions are bound along with ATP. From these observations we surmise that interaction of basic residues with ATP in P2X2 are critical for robust opening of the channel and the presence of a divalent ion bound to ATP weakens those interactions. For example, switching of a salt bridge between the equivalent of R302 and E179 to between R302 and ATP has been proposed to promote opening of rP2X2 (*49*) and binding of Mg^2+^ to ATP might prevent that switch. To look for potential interactions of Mg^2+^ with P2X2R in the absence of ATP we also determined a structure of hP2X2-S in the presence of Mg^2+^ (5 mM) alone at a resolution of 3 Å (**Fig. S3; Table S2**), which was indistinguishable from the structure of apo hP2X2-S with no observable density for Mg^2+^ (**Fig. S4b**).

We next investigated how the antagonist suramin binds to P2X2Rs. Suramin is one of the earliest antagonists used to study P2XRs (*50*), many derivatives with improved selectivity for P2XRs have been synthesized (*51*) and it functions as a partial agonist in rP2X2 when the closed gate is destabilized by the equivalent of the T350S mutation (*52*). We succeeded in solving a structure of hP2X2-L with suramin at a resolution of 3.2 Å (**Fig. 5a-d; Fig. S7; Table S1**). Suramin adopts a symmetric anchor shape, with the blades of the anchor sinking deep into the protein and partially overlapping with the ATP site (**Fig. 5a-d**), very different than predicted by docking (*51*). One blade inserts between the DF of one subunit and the LF of an adjacent subunit, and the other into a pocket between the same DF and the HD of the adjacent subunit (**Fig. 5c,d**). By contrast, ATP binds to an asymmetric pocket formed by two adjacent subunits, interacting with the HD, UB and LF of one subunit and the LB and DF of the adjacent subunit (**Fig. 2d**). Suramin would prevent ATP from engaging with the receptor, suggesting that it is a competitive antagonist, in contrast to the proposal that it binds to a distinct site from ATP and acts as an allosteric inhibitor (*52*). We do observe modest conformational changes in the HD, DF and LF (**Fig. 5d**), which may explain how suramin can act as a partial agonist when the closed state is destabilized (*52*), yet in our structure the activation gate remains fully closed (**Fig. 5e; Fig. S8e**).

**Fig. 5.**
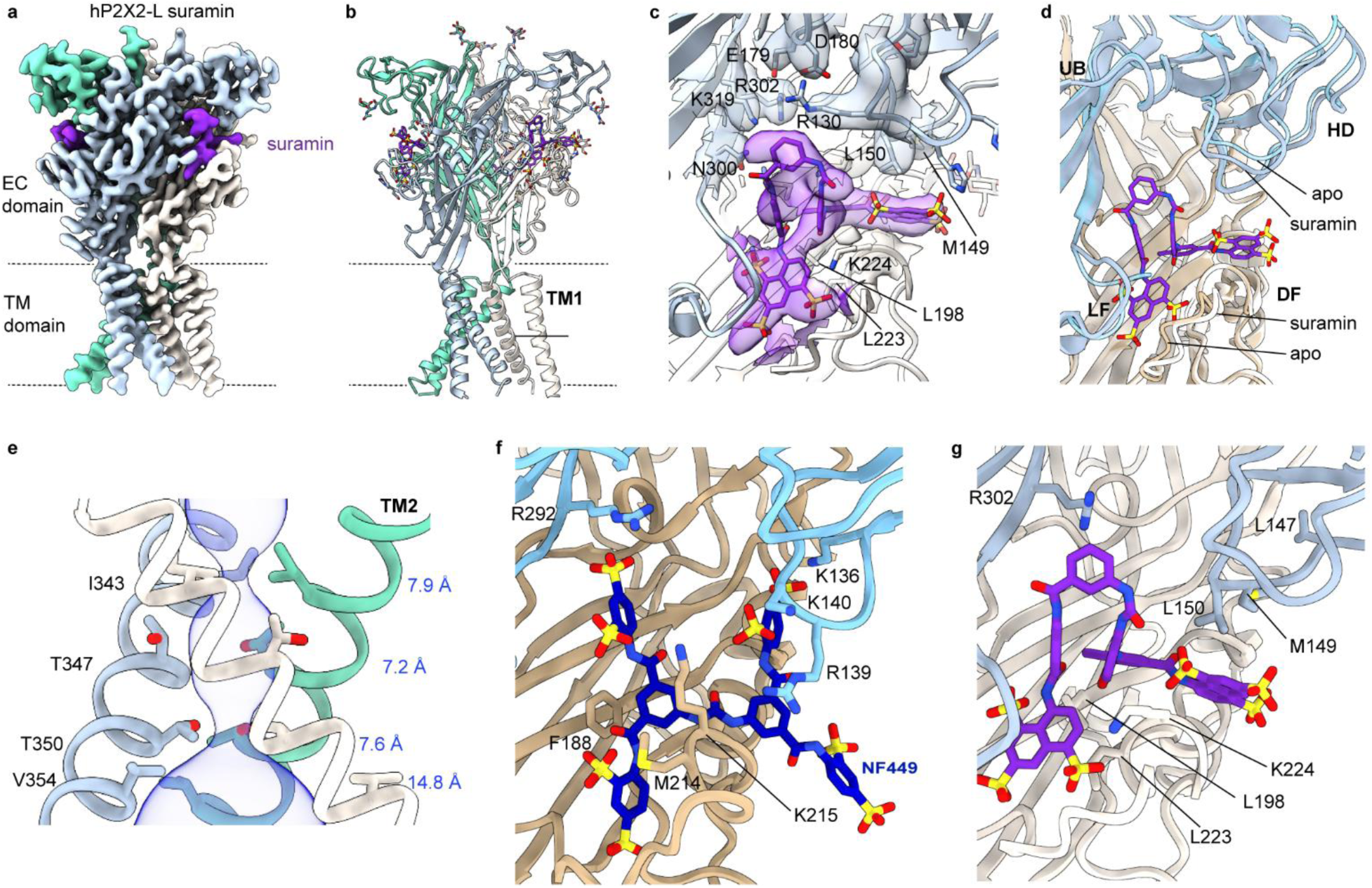
Structure of the hP2X2R with suramin bound. **a**) Cryo-EM map for hP2X2-L with suramin bound (DeepEMhancer sharpened). Density for suramin is shown in violet. **b**) Structural model for hP2X2-L with suramin bound. **c**) Structure of hP2X2-L with suramin bound within the agonist recognition pocket. Cryo-EM density is shown for key side chains near to where suramin binds (DeepEMhancer sharpened, contour ∼0.05). **d**) Similar view as in c but superimposing structures of apo and suramin bound hP2X2-L. **e**) Structure of the activation gate region for hP2X2-L with suramin bound shown as a side view along with MOLE (*73*) representation. Residues in the regions of the pore that function as gates are shown as stick representation with distances shown in blue between adjacent Cα atoms for I343, T347, T350 and V354. **f**) Structural model of hP2X1 with NF449 bound (9b95) shown in stick representation with residues implicated in NF449 inhibition shown (see text). **g**) Structural model of hP2X2 with suramin in a similar orientation as in panel f with residues corresponding to those in panel f also shown.

We also compared our suramin bound structure with the highest resolution structure available for P2X1 bound by NF449 (*21*), a suramin derivative that has 1000-fold selectivity for hP2X1 compared to hP2X2 (*53*). NF449 inserts into the ATP binding pocket and like suramin extends into one pocket between the DF of one subunit and the LF of an adjacent subunit, and another between the same DF and the HD of the adjacent subunit (**Fig. 5f,g**). NF449 has an additional sulfonic acid group that reaches up to R292 in the UB of hP2X1 (**Fig. 5f**), which is essential for activation by ATP and inhibition by NF449 (*20, 21*), whereas suramin lacks functional groups to interact with the equivalent R302 residue in hP2X2 (**Fig. 5g**), helping to explain the higher affinity of NF449 for both hP2X1 and hP2X2 compared to suramin (*53*). NF449 also contacts K136, R139 and K140 in the HD of hP2X1 (**Fig. 5f**), positions where mutations greatly diminish inhibition by the antagonist (*20, 21, 53*). In contrast, the equivalent residues in the HD of hP2X2 are L147, M149 and L150, and only L150 makes a hydrophobic interaction with suramin (**Fig. 5g**), suggesting that these residues in the HD are more important determinants for NF449 binding to hP2X1 than suramin binding to hP2X2. M214 and K215 in the DF of hP2X1 also contact NF449 (**Fig. 5f**) and their mutation diminishes inhibition by the antagonist (*21*), and the corresponding residues in hP2X2 (L223 and K224) are conserved and contact suramin in our structure (**Fig. 5g**). These comparisons suggest that the structure of suramin bound to hP2X2 provides a foundation for development of the suramin scaffold to improve selectivity towards hP2X2. For example, modification of suramin to interact with the R130, L147 and M149 in the HD would be expected to enhance selectivity towards hP2X2 as these residues are not conserved in other P2XRs (**Fig. S2**).

### Mechanism of channel opening in a membrane-like environment

Comparison of the structures of apo hP2X2-L in nanodiscs with the four classes bound by ATP^4-^ reveals that the nucleotide triggers distinct conformational changes in the TMD. For class 1 (∼11 % of particles), the activation gate expands at the extracellular end of the pore at I343 while remaining narrow at T350 (**Fig. 6a; Fig. S8**). For class 2 (∼18 % of particles), the entire activation gate widens sufficiently for hydrated cations to permeate, with Oγ of T347 and T350 positioned where they would interact with permeating ions (**Fig. 6b,e,f; Fig. S8**). For class 3 (∼26 % of particles), the external end of the activation gate remains open but the internal pore near V354 begins to narrow (**Fig. 6c; Fig. S8**). Similarly for class 4 (∼7 % of particles), the activation gate remains open while the internal pore becomes occluded by V354 (**Fig. 6d,g,h; Fig. S8**).

**Fig. 6.**
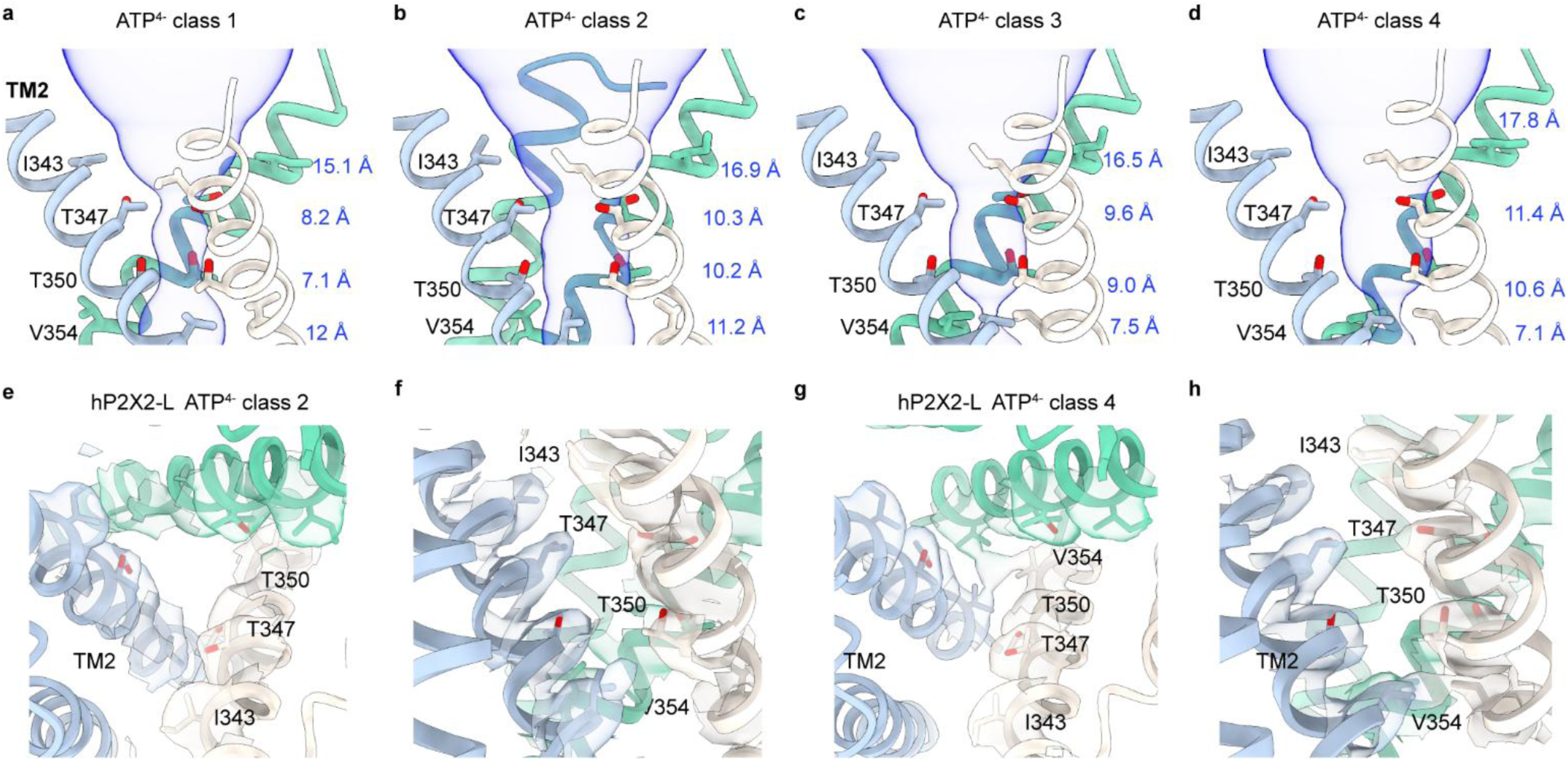
Conformation changes within the TM2 helix of hP2X2 upon ATP^4-^ binding. **a-d**) MOLE (*73*) representations for the pore of the four classes with ATP^4-^ bound. Residues in the regions of the pore that function as gates are shown as stick representation with distances shown between adjacent Cα for I343, T347, T350 and V354. **e,f**) Structure of the activation gate in the pore of ATP^4-^ bound hP2X2-L class 2 viewed from the extracellular side of the membrane (e) or as a side view (f). Residues in the gate region are shown as stick representation with cryo-EM density for side chains (DeepEMhancer sharpened, contour ∼0.07). **g,h**) Structure of the activation gate in the pore of ATP^4-^ bound hP2X2-L class 4 viewed from the extracellular side of the membrane (g) or as a side view (h). Residues in the gate region are shown as stick representation with cryo-EM density for side chains (DeepEMhancer sharpened, contour ∼0.09).

Although assignment of conformations seen in cryo-EM structures to functional states is always challenging, we think that we can reasonably assign two of these classes to distinct functional states. As discussed above, cysteine accessibility studies support the location of an activation gate involving I343, T347 and T350 (*26, 27*), which in apo hP2X2-L form a constriction in the pore and thus the apo structure likely represents a closed state (**Fig. 1d**). Class 2 in the presence of ATP^4-^ likely represents an open state as the residues that form the gate in the closed state expand and become exposed to solvent, with radii varying between 5.4 Å at N344 and 3.2 Å at T350 (**Fig. 6b,e,f**), consistent with accessibility data demonstrating that Cys substitutions within the gate react rapidly with thiol reactive compounds or Ag^+^ ions in the open state (*26, 27*). The conformational changes occurring between apo and open states involve translation of the TM2 helix by one helical turn (∼6 Å displacement of L338 Cα) towards the extracellular side and rotation of ∼16°, together leading to a dramatic expansion of the activation gate to open the channel (**Fig. 6b,e; Fig. S8b,e; Movie S1**). The radius of the activation gate at its narrowest point is ∼3 Å, sufficient to support the permeation of hydrated Na^+^ and Ca^2+^ ions.

Although we did not anticipate capturing hP2X2 in a desensitized state given that P2X2 desensitizes on the timescale of secs and we used brief exposure to ATP^4-^, it seems likely that class 4 represents a desensitized state because the activation gate remains open (3.3 to 5.4 Å radii at N344, T347 and T350), but the pore narrows at V354 to a radius of ∼0.6 Å (**Fig.6d,g,h**). Class 4 representing a desensitized state would be consistent with the desensitized states proposed for P2X3 determined with endogenous ATP bound (*17*) and more recently for hP2X1, hP2X2 and hP2X4 obtained using similar approaches (*17, 20–23*) (**Fig. S9c,h-k**), suggesting that the mechanisms of desensitization are relatively conserved in P2XRs even though the kinetics of desensitization vary widely. Overall, our assignment of closed, open and desensitized states for hP2X2Rs in nanodiscs are similar to those determined for hP2X3Rs in detergents (*17*). The two remaining classes (class 1 and 3) likely represent intermediate states that have not previously been described for P2XRs. The structures we determined for hP2X2 in detergent with residual ATP bound superimpose well with class 1 with ATP^4-^ (**Fig. S9d-g**), suggesting that all three structures represent a similar intermediate state. One possibility is that class 1 represents a pre-open state and class 3 represents a pre-desensitize state as a morph between apo to class 1 through 4 shows gradual transitions from closed to open to desensitized states (**Movie S1**).

In the available structures of P2X3, P2X4 and P2X7 captured in open states, the subunit interfaces in membrane spanning regions completely sever as the channel expands during opening (*15–19*). This feature of those open state structures is evident in side views where adjacent subunits no longer interact (**Fig. 7a-c**), which in simulations allow lipids to diffuse into the pore to interfere with ion permeation (*24*). For the open states of P2X3 and P2X7, the buried surface area between neighboring TM2 helices was 65 Å^2^ and 200 Å^2^, respectively, but most or all of this buried surface is attributable to the cytoplasmic caps for P2X3 (57 Å^2^) and P2X7 (200 Å^2^). In contrast, for structures of hP2X2 with ATP^4-^ bound, these subunit interfaces remain intact in all four classes, with buried surface area ranging from 130 Å^2^ for class 2 to 161 Å^2^ for class 3 (**Fig. 7d,e and legend**). For class 2 of hP2X2, the subunit interface stabilizing the open state is formed by L349, T350, S351, V352, G353, V354, S356, F357 and W361 (**Fig. 7d,e**). It is notable that T350 within the activation gate contributes to the subunit interface in the outer leaflet, which is possible because TM2 rotates away from the central axis during channel opening. Although a subunit interface is maintained as hP2X2 opens, we see a lateral portal within the membrane, a feature seen in other ion channels where they have been hypothesized to enable access of hydrophobic drugs to the pore (*54–56*). We do not see density corresponding to lipids inside the pore in the open state even though density for lipids is evident outside the TM helices (**Fig. S8f,g**), indicating that lipids cannot enter the pore but help to stabilize subunit packing. We do see density for several lipid molecules outside the pore in the open state, although they are not as well-resolved as those seen in the closed state (**Fig. 1l**).

**Fig. 7.**
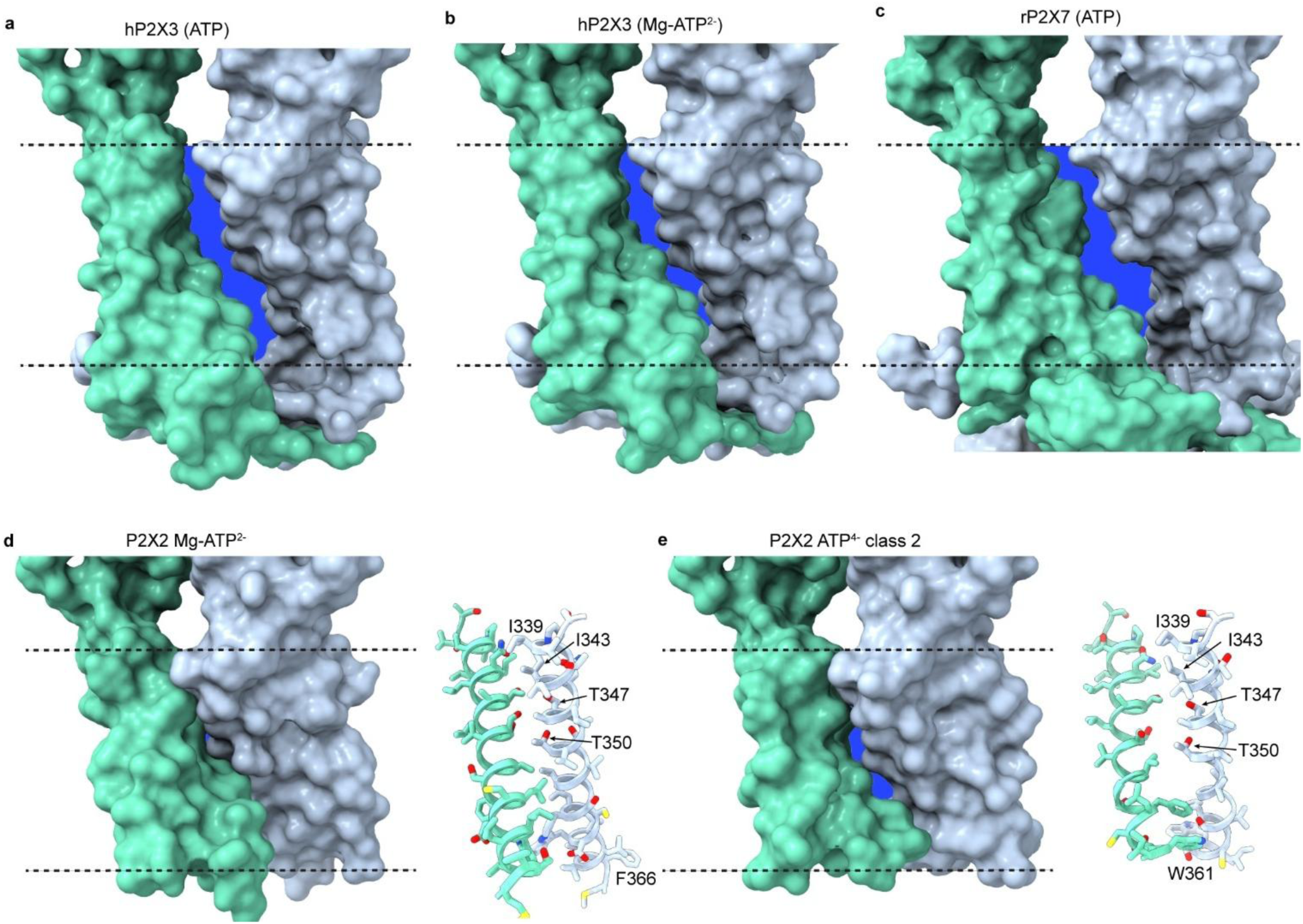
Subunit interfaces within the TM domain are retained for hP2X2 in nanodiscs. **a**) Surface rendering of the structure of the open state of hP2X3 with ATP bound (5svk). **b**) Surface rendering of the structure of the open state of hP2X3 with Mg-ATP^2-^ bound (6ah4). **c**) Surface rendering of the structure of the open state of rP2X7 with ATP bound (6u9w). **d**) Surface representation of the TM region and side chain interactions between adjacent TM2 helices for Mg-ATP^2-^ bound to hP2X2-S. **e**) Surface representation of the TM region and side chain interactions between adjacent TM2 helices for class 2 with ATP^4-^ bound to hP2X2-L. For all panels, the subunit interfaces within the TMD are shown on a blue background to illustrate gaps between subunits. Solvent accessible surface area and buried surface area, respectively, between the TM2 helices lining the pore region of chains A and B were 4318 Å^2^ and 65 Å^2^ for hP2X3 with ATP, 4560 Å^2^ and 190 Å^2^ for hP2X3 with Mg-ATP^2-^ (2628 Å^2^ and 252 Å^2^ for hP2X3 apo), 5934 Å^2^ and 200 Å^2^ for rP2X7 with ATP (5669 Å^2^ and 402 Å^2^ for rP2X7 apo), 3014 Å^2^ and 186 Å^2^ for hP2X2-S with Mg-ATP^2-^, 2428 Å^2^ and 130 Å^2^ for hP2X2-L with ATP^4-^ (2945 Å^2^ and 210 Å^2^ for hP2X2-L apo). Surface area was calculated using PDB-ePISA, http://pdbe.org/pisa/).

## Discussion

The goals of this study were to determine structures of hP2X2R to understand the unique functional properties of this subtype, and to explore how the membrane lipids influence the structure and dynamics of the protein. One of the striking features of P2X2Rs is that they require ATP^4-^ for activation (*30, 42*). In physiological solutions containing mM concentrations of Mg^2+^ and Ca^2+^, the most prevalent forms of ATP will be bound by divalent ions. While Mg-ATP^2-^ and Ca-ATP^2-^ can bind to P2X2R, they are ineffective at promoting channel opening (*30, 42*) (**Fig. 3**). In contrast, in hP2X3Rs an extended acidic chamber comprised of four residues plays a critical role in enabling activation by ATP bound by divalent ions (*19*). The acidic chamber in hP2X3Rs can bind a divalent ion in the absence of ATP (*17*) and when ATP binds that divalent ion interacts with the γ-PO_4_ of ATP on the receptor and catalyzes the release of the divalent ion that binds along with ATP, effectively extracting ATP^4-^ from a physiological solution containing mostly ATP bound by divalent ions (*19*). Our structures of hP2X2R reveal that this subtype contains a unique Arg residue (R130) that interacts with the γ-PO_4_ of ATP (**Fig. 2**), effectively replacing the divalent ion that stabilizes ATP in P2X3Rs (*19*). The position of two acidic residues that are present hP2X2Rs in the vicinity of ATP appear to be suboptimal for supporting activation by Mg-ATP^2-^because the R130A mutant could not be activated by Mg-ATP^2-^ (**Fig. 3**). However, we found that the R130E mutation was sufficient to enable activation of hP2X2R by both ATP^4-^ and Mg-ATP^2-^(**Fig. 3**), indicating that an acidic residue at position 130 in hP2X2 receptors can interact with Mg-ATP^2-^ to open the channel. In structures solved in the presence of Mg-ATP^2-^ we see conformational changes in the ECD and outer ends of TM2 that are intermediate between apo and ATP^4-^ bound forms of the receptor while the lower part of the activation gate remains closed (**Fig. 4**), indicating that Mg-ATP^2-^ is bound (**Fig. 3**). In addition, the structure of suramin bound to hP2X2 demonstrate that this inhibitor is a competitive antagonist, and all of our structures provide a blueprint for developing therapeutics that selectively target this receptor subtype.

Having structures of hP2X2Rs in closed, open and with Mg-ATP^2-^ provides an opportunity to examine structural elements that may be involved in coupling ATP^4-^ binding to channel opening. Within the ECD near the membrane, intersubunit interaction between the β10-β11 loop and the β14 strand appear to stabilize the closed state (**Fig. 8a**). H330 in β14 of one subunit interacts with P206 in the adjacent subunit in the closed state and in structures with either ATP^4-^ or Mg-ATP^2-^ this interaction breaks as H330 swings upward to interact with E75 in β2 (**Fig. 8a-c; Movie S2**). The β14 strand where H330 is located directly connects to the pore-lining TM2 helix where the gate resides and P206 is in the β10-β11 loop where the β10, β11 and nearby β2 strands extend up into the ATP binding pocket (**Fig. 8a-c; Fig. S10; Movie S2**), providing a direct pathway for ATP binding to tug on β2, β10 and β11 to weaken the interactions of P206 and H330. In P2X2Rs, H330 (H319 in rP2X2R) has been implicated in proton potentiation (*57, 58*) and P206 and the surrounding His residues have been implicated in the regulation by Zn^2+^ (*58, 59*), which in hP2X2 inhibits opening of the channel. However, the role of these residues in regulation of the channel by Zn^2+^ and protons will need to be reexamined because previous studies were done in the presence of divalent cations (*57–59*) and the requirement for ATP^4-^ was not yet appreciated (*30*). Indeed, lowering pH will increase ATP^4-^ concentrations in the presence of divalent cations and Zn^2+^ binds ATP^4-^ more tightly than either Ca^2+^ or Mg^2+^ and will therefore decrease ATP^4-^ concentrations in physiological solutions. However, the H319K mutation in rP2X2R increases the apparent affinity for ATP by approximately 30-fold (*57, 60*), consistent with the structural changes we see between closed, ATP^4-^ and Mg-ATP^2-^ bound hP2XRs (**Fig. 8a-c**). Just below H330 and P206, T72 residues in the β1 strands are positioned at the subunit interface where their interaction would stabilize the closed state (4.3 Å, Cγ2 to Cγ2) (**Fig. 8a**). In our structures with ATP^4-^ bound we see two conformations of the backbone between T72 and P74 in β1 where T72 moves away from the subunit interface to different extents (8.3 Å, Cγ2 to Cγ2 for conformation 1 and 10.5 Å, Cγ2 to Cγ2 for conformation 2) (**Fig. S5e**), suggesting that this local region is conformationally heterogeneous. In our structure with Mg-ATP^2-^ bound, most of the β1 strand has moved outward as the β-strand network expands, but a distinctive bend in the β1 strand is produced by T72 residues remaining positioned closer together (6.8 Å, Cγ2 to Cγ2) (**Fig. 8b**). These structural observations suggest that the intersubunit interaction of H330 with P206 and those between T72 residues likely serve to stabilize the ECD in a closed conformation and that releasing these interactions is necessary but not sufficient to open the channel.

**Fig. 8.**
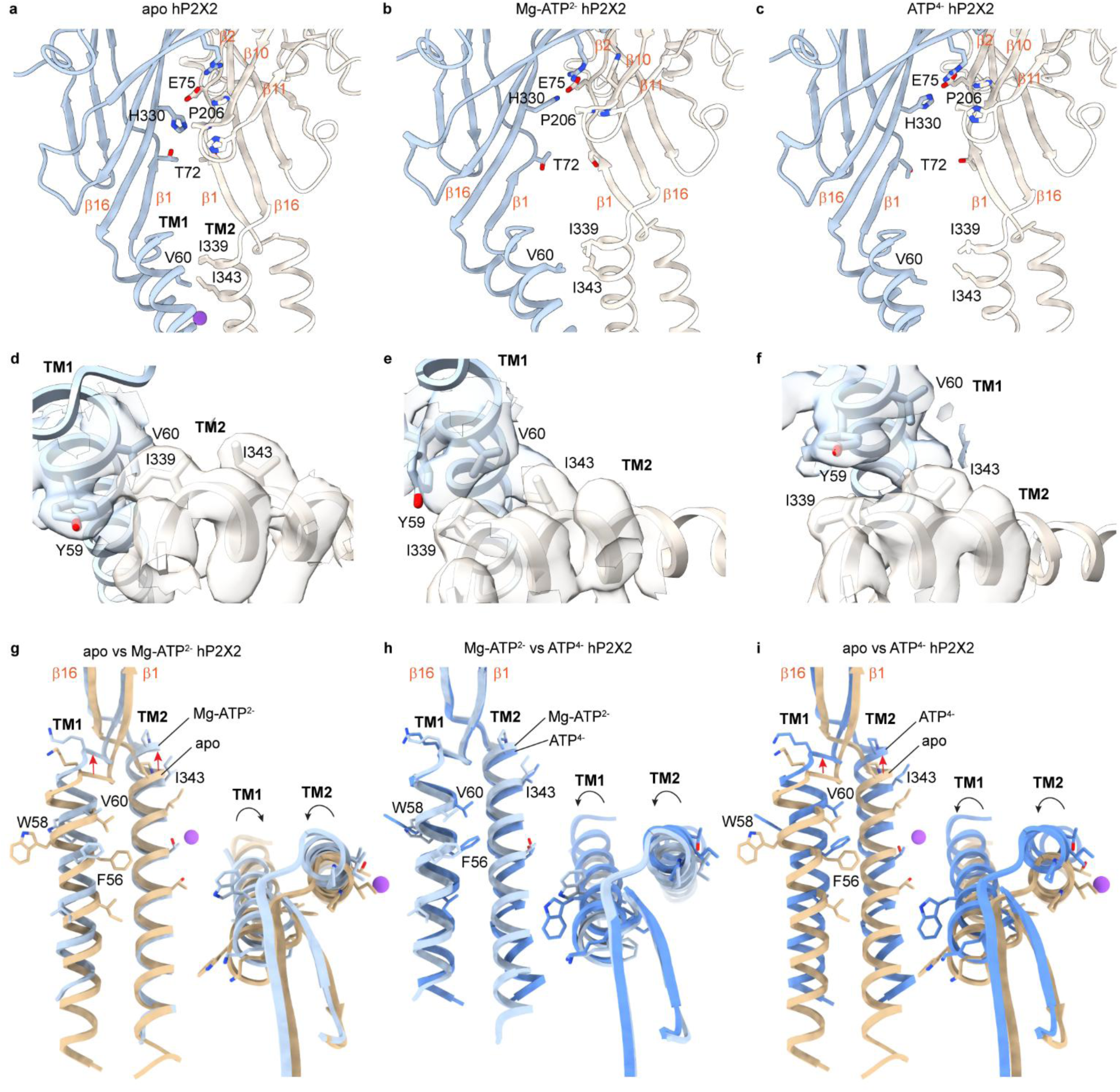
Regulatory elements involved in coupling ATP binding to channel opening. **a-c**) Structural models for apo hP2X2-L, Mg-ATP^2-^ hP2X2-S and ATP^4-^ hP2X2-L (class 2), respectively, highlighting the location of regulatory elements involved in coupling ATP binding to channel opening. **d-f**) Cryo-EM density and structural model for apo hP2X2-L, Mg-ATP^2-^hP2X2-S and ATP^4-^ hP2X2-L (class 2), respectively, showing dynamic interactions between V60 in TM1 and I339 in TM2 in adjacent subunits. Cryo-EM maps were sharpened using DeepEMhancer (contour ∼0.04, 0.1, 0.05 respectively). **g-i**) Superimposed TM helices from a single subunit for apo and Mg-ATP^2-^ (**g**), Mg-ATP^2-^ and ATP^4-^ (**h**) and apo and ATP^4-^ (**i**) illustrating translations and rotations of the two TMs between the three conditions. Purple sphere is a Na^+^ ion modeled for apo. Insets for panels g-i are views from the extracellular side.

A critical determinant governing channel opening within the TM region is the intersubunit interaction between V60 and I339 that is known to be critical for stabilizing the closed state. The V60L hearing loss mutation in hP2X2 causes constitutive activation of the channel and uncoupling of ATP binding and channel opening (*36*), indicating that even subtle structural changes at this position have a profound impact on activation of the channel. In addition, disulfide bonds between Cys substitutions at residues equivalent to V60 (V48) and I339 (I328) lock rP2X2R closed (*37, 38*) and rP2X2Rs containing the equivalent of I339C can be directly activated by lipophilic thiol reactive compounds (*39*), indicating that the interaction of these two residues plays a particularly important role in stabilizing the closed state and sensing the binding of ATP. In the closed state, V60 in TM1 and I339 in TM2 of the adjacent subunit exhibit hydrophobic interactions at the subunit interface (4.5 Å, Cγ2 to Cγ2) (**Fig. 8a,d**) and with partially resolved lipids nearby (**Fig. 1l**). Interaction between V60 and I339 are retained when Mg-ATP^2-^ binds (4.2 Å, Cγ2 to Cγ2) (**Fig. 8b,e**) because as the TM2 helix moves outward and begins to rotate in a counterclockwise direction viewed from the outside, the TM1 helix rotates in a clockwise direction as it also moves outward (**Fig. 8g,h; Fig. S8h**). In contrast, when ATP^4-^ binds, TM1 and TM2 fully rotate, both in counterclockwise directions when viewed from the outside (**Fig. 8i; Fig. S8h**), breaking the interaction of V60 and I339 (9.54 Å, Cγ1 to Cγ2) as the activation gate opens (**Fig. 8c,f**). Although the resolution of our structures is insufficient to pinpoint why Mg-ATP^2-^ binding is not able to open the channel, it seems likely that relatively subtle structural changes in β1 strands are needed for the directly connected TM1 helix to fully rotate and break its interaction with I339. The distinct rotations of the TM1 helix seen when comparing apo, Mg-ATP^2-^ and ATP^4-^ suggest a critical role of this peripheral TM1 helix in channel opening, consistent with the pronounced influence that mutations in TM1 have on gating of P2X2Rs (*38, 61*).

One technical challenge in determining structures of P2XRs in different conformational states is depleting the ATP binding site of endogenous ATP that copurifies with these proteins. We succeeded in depleting ATP for the protein reconstituted into nanodiscs using apyrase and alkaline phosphatase, but this manipulation failed to completely deplete endogenous ATP when hP2XR was purified in DDM (**Fig. S4c**), suggesting that DDM may restrict the conformation of P2XRs. Lipid nanodiscs do provide a more membrane-like environment, perhaps allowing the protein to explore more conformational states and release endogenous ATP. Indeed, a recent study reporting the structure of apo hP2X2 also succeeded in removing ATP using apyrase in DDM along with cholesteryl hemisuccinate (*22*), which would more closely mimic a native membrane environment.

Our structures of P2X2Rs in lipid nanodiscs also indicate that the lipid membrane plays a critical role in influencing the structure and dynamics of the TM region of these receptors. The influence of membrane lipids helps to explain why we observe so many distinct conformations in lipid nanodiscs (**Fig. 6**), including intermediate states not previously reported. We also see extensive interactions of lipids with the apo state of P2X2Rs in the outer leaflet of the bilayer where the membrane embraces the TMD (**Fig. 1l**), suggesting that the membrane could help to stabilize the closed state of the channel. Unlike previous structures in detergents where the interactions between adjacent subunits break as the channel opens, the structures in nanodiscs retain intact subunit interfaces (**Fig. 7d,e**), suggesting that the membrane constrains the TM region during opening. Although density for the cytoplasmic cap resolved in other subtypes (*17, 18, 23*) was not sufficiently resolved to model this region in P2X2Rs, the density we do see suggests that it is likely present in all states of the gating cycle. In the future it will be interesting to determine structures of P2XRs in other membrane mimetics or in lipid vesicles to understand how the membrane deforms around the internal ends of the TMs where lateral fenestrations provide a pathway for ions to enter and exit the internal pore (*44*).

## Materials and Methods

### hP2X2R expression using Baculovirus and mammalian expression system

To produce the hP2X2R for cryo-EM, the codon optimized genes of both the splice variants (hP2X2-L, *Q9UBL9-1* and hP2X2-S, *Q9UBL9-2*) were cloned into the pEGBacMam vector with the (His)_8_- affinity and mVenus fluorescence tag (*62*) on the N-terminus and a TEV protease site between the fluorescence tag and the receptor. These constructs were expressed in tsA201 cells using the previously published Baculovirus-mammalian expression system with a few minor modifications (*63*). Briefly, bacmids were purified from the transformed DH10Bac *E. coli* cells using a standard alkaline lysis method. P0 virus was generated by transfecting Sf9 cells (∼0.5×10^6^ cells/mL in a T25 flask) with 1 µg/mL of fresh bacmid using Cellfectin and incubated at 27 °C for 7 days. To amplify the P0 virus, ∼500 mL of suspension culture of Sf9 cells at ∼0.5×10^6^ cells/mL were infected with 500-750 μL of the P0 virus and incubated at 27 °C for 5 days. The P1 virus was harvested by centrifugation (4,781 g x 45 min, 4 °C), filtered through 0.22 μm filter, supplemented with 1 % FBS and stored in dark at 4 °C. To obtain the hP2X2 proteins, 2.4 L of tsA201 cells (∼1.5×10^6^ cells/mL) in Freestyle media with 1 % FBS and 0.25x Penstrep (Gibco, 10,000 U/mL) were infected with 4-6 % (v/v) of P1 virus and incubated at 37 °C with 8% CO_2_. To boost protein production, 10 mM sodium butyrate was added at ∼16 hrs in the infected cells and the cultures were shifted to 30 °C, 8% CO_2_ for 48 hrs post-infection. The cells were harvested by centrifugation (4,781g x 45 min, 4 °C), washed with ice-cold 1X TBS (20 mM Tris pH 7.4 and 150 mM NaCl) and stored at −80 °C until further use.

### hP2X2R purification

To purify the hP2X2R, the frozen cells were thawed and then resuspended in ice-cold 50 mM Tris pH 8, 150 mM NaCl, supplemented with a cOmplete^TM^ Protease Inhibitor Cocktail Tablet (1 tablet/ 50 mL lysate) and 1 mM PMSF. Cells were lysed using a Dounce homogenizer followed by sonication with QSonica Q700 sonicator in three/four rounds of 90 seconds each, at a power level of 30 (Pulse ON: 10 s, Pulse OFF: 20 s, temperature cut-off: 12 °C). The lysate was clarified by low-speed centrifugation at 10,480 g for 20 min to remove cell debris and intact cells. The membranes were obtained by ultracentrifugation at 125,000 g for 1.25 hr at 4 °C, flash frozen and stored at −80 °C until further use. Our initial attempts to obtain the apo receptor were hampered by the presence of residual ATP in the hP2X2 binding pocket derived from lysed cells and retained throughout the purification. Hence, we added 1 U/mL each of apyrase (A6410, Sigma) and alkaline phosphatase (M0371L, NEB) during the cell lysis and membrane solubilization step to remove any residual ATP. To purify the hP2X2Rs, the membranes were resuspended in equilibration buffer (50 mM Tris pH 8, 150 mM NaCl, 1 mM PMSF, 2 mM MgCl_2_, 1 U/mL apyrase, 1 U/mL alkaline phosphatase, 10 mM imidazole and cOmplete^TM^ Protease Inhibitor Cocktail Tablet), homogenized using Dounce homogenizer and solubilized with 30 mM DDM for 1 hr at 4 °C on a rotator. The lysate was clarified by centrifugation (43,065 g for 30 min, 4 °C) and incubated with pre-equilibrated CoTALON resin (Clontech, 2 mL resin/ L culture) at 4 °C for 2.5 hrs. Post incubation, the column was packed, and the unbound fraction was passed through the packed resin twice. The resin was washed with 10 column volume (CV) of equilibration buffer followed by 50 CV of wash buffer (50 mM Tris pH 8.2, 150 mM NaCl, 2 mM MgCl_2_, 0.5 mM DDM, and 40 mM imidazole). The protein was eluted in 0.5x CV fractions with elution buffer (50 mM Tris pH 8, 150 mM NaCl, 0.5 mM DDM, and 300 mM imidazole) until OD_280 nm_ ≤0. The eluate fractions ≥1 mg/mL were directly used for nanodisc reconstitution and rest of the fractions (0.3-0.9 mg/mL) were pooled and concentrated up to 1-1.2 mg/mL using Amicon Ultra (100 kDa cutoff) before nanodisc reconstitution. The protein was incubated overnight with TEV (prepared in-house) and 2 mM TCEP for removal of the fluorescent/affinity tags prior to nanodisc reconstitution. All purification steps described above were carried out at 4 °C or on ice.

### Lipid nanodisc reconstitution of the hP2X2R

Lipid nanodisc reconstitution was performed following the previously published methods with minor modifications (*64, 65*). The membrane scaffold protein MSP1E3D1 was expressed and purified using pMSP1E3D1 (Addgene #20066) in BL21-CodonPlus(DE3)-RIL *E coli* cells (Agilent) according to a published protocol (*65*). For nanodisc reconstitution, the purified hP2X2R in detergent was incubated with MSP1E3D1, 3:1:1 mixture of 1-palmitoyl-2-oleoyl-sn-glycero-3-phosphocholine (POPC), 1-palmitoyl-2-oleoyl-sn-glycero-3-phospho-(1’-rac-glycerol) (POPG) and 1-palmitoyl-2-oleoyl-sn-glycero-3-phosphoethanolamine (POPE), with or without porcine brain PIP2 (Avanti Polar Lipids), added as 25 % of total lipids and 14 mM sodium cholate for 30-45 min, at room temperature. The reconstitution was done at 1 (P2X2):2 (MSP):60 (lipids) molar ratio. The mixture was transferred to a tube with SM-2 Biobeads (∼30-50 fold of detergent; w/w) and incubated at room temperature for ∼2 hrs with rotation and the Biobeads were replaced each hr. The reconstituted protein was concentrated using Amicon Ultra (100 kDa cutoff), and separated from aggregates, empty nanodiscs and cleaved tags using size exclusion chromatography (Superose 6 increase column, 10/300 GL) in a buffer containing 20 mM HEPES pH 7, 100 mM NaCl. The peak fractions were run on SDS-PAGE (stain-free, Biorad) to verify the presence of the receptor and MSP1E3D1, fractions were pooled and concentrated up to ∼4.5 mg/mL.

### Cryo-EM sample preparation and data acquisition

The purified hP2X2 protein samples in nanodisc or detergent (2.5-3 µL) with freshly added 0.5 −0.75 mM fluorinated Fos-Choline-8 (Anatrace) were applied to glow discharged Ultrafoil or Quantifoil grids (R 1.2/1.3 Au, 300 mesh). For samples with ATP^4-^ and Mg-ATP^2-^, the agonists were applied to the protein on the grid immediately before blotting for 2.5 s. For the suramin (0.5 mM from a 3 mM stock in H_2_O) dataset, the compound was added to the protein 1-2 h prior to the grid preparation. The grids were blotted for 2.5 or 3 s, blot-force 0, 100% humidity, at 16 °C using a FEI Vitrobot Mark IV (Thermo Fisher), followed by plunging into liquid ethane cooled by liquid nitrogen.

Images were acquired using an FEI Titan Krios equipped with a Gatan LS image energy filter (slit width 20 eV) operating at 300 kV. A Gatan K3 Summit direct electron detector was used to record movies in super-resolution mode with a nominal magnification of 105,000 x, resulting in a calibrated pixel size of 0.415 or 0.412 Å per pixel. The typical defocus values ranged from −0.8 to −2.4 um. Exposures of 2.2 s were dose-fractionated into 40 frames, resulting in a total dose of ∼60 e^-^ Å^-2^. Movies were recorded using the automated acquisition program SerialEM (*66*) or EPU v3.3.1 (Thermo Scientific).

### Image processing

All datasets were processed in cryoSPARC v4.5.3 (*67*). In brief, the beam-induced motion between frames of each movie was corrected and Fourier binned by 2 using Patch Motion Correction and contrast transfer function (CTF) estimation was performed using Patch CTF estimation. Micrographs were manually curated based on CTF fit estimation values (cut-off: ∼5-6 Å). Automated particle picking was done from a small subset of micrographs using Blob picker, followed by reference-free 2D Classification. Ab-initio reconstruction and heterogenous classification was then done to remove junk particles. The good class was then refined using homogenous and non-uniform refinement followed by template generation. This template was used to pick particles from manually curated micrographs using template picker. The particles were inspected to estimate thresholds for particle extraction followed by removal of duplicate particles. Best particles were selected using 3D classification that included *ab-initio* and heterogenous classification steps iteratively. The best class(es) was/were refined using non-uniform refinement with C1 or C3 symmetry followed by reference-based motion correction. For structures of apo hP2X2, the particles were further cleaned up by a final round of heterogenous classification. CTF refinement was done prior to local refinement with C3 symmetry to obtain the final map. For ATP^4-^ and Mg-ATP^2-^datasets, another round of 3D classification was done using a focused mask around the TM/cytoplasmic cap region or ATP binding pocket respectively to capture all possible classes. Each class was individually refined to obtain high-resolution maps and then compared to obtain different states with ATP^4-^ or with a uniform density of ATP in all subunits for Mg-ATP^2-^. The resolutions of final maps for all datasets in this study were between 2.6 and 3.2 Å. The maps were sharpened in cryoSPARC or deepEMhancer (*68*) (with tightTarget/ highRes models) and used for model building.

### Model building and structure refinement

AlphaFold 3 (*69*) was used to build initial models for hP2X2-L and hP2X2-S apo states and ATP^4-^ or Mg-ATP^2-^ bound states. These were docked into the respective maps using UCSF ChimeraX v1.10 (*70*) and the model for each structure was refined using PHENIX v1.20.1 (*71*) with secondary structure and geometry restraints. The outliers were manually corrected in WinCoot v0.9.8.95 (*72*) to fit the density. The final models were evaluated by comprehensive validation in PHENIX and wwPDB validation system (https://validate.wwpdb.org). Structural visualizations were generated using UCSF ChimeraX and WinCoot. Contour levels provided in figure legends are from ChimeraX. MOLE representations were generated using MOLEonline 2.5 (https://moleonline.cz/) (*73*). Surface accessibility was calculated using PDB-ePISA server (http://pdbe.org/pisa/). The Na^+^ bound within the activation gate of apo hP2X2 was evaluated using CheckMyMetal (http://cmm.minorlab.org).

### Structure-based and sequence-based alignments

Structure-based sequence alignments of different apo P2XRs to apo hP2X2-L was performed using PROMALS3D (*74*) and sequence-based alignment for the two splice variants of hP2X2 was performed using Clustal Omega (*75*). Both the alignments were visualized using ESPript 3.0 (*76*) with a %Equivalent coloring scheme.

### Electrophysiological Recordings

WT hP2X2-L, R130A and R130E mutants were cloned in IRES GFP-pcDNA3.1 or pcDNA3.1. Accutase-treated HEK293 cells were seeded onto glass coverslips in six-well plates in DMEM supplemented with 10% (v/v) FBS and 10 mg/L gentamicin 4-6 h before transfection and placed in a 37 °C incubator with 5 % CO_2_. 0.5-0.75 µg WT hP2X2L or R130A IRES GFP-pcDNA3.1 construct or R130E pcDNA3.1 (co-transfected with GFP cDNA in pGreen-Lantern; ratio of 10:1) was transfected using FuGENE6 Transfection Reagent (Roche Applied Science). Cells were used for whole-cell recording 16-26 h post transfection. Whole-cell patch-clamp recording was used to record hP2X2R channel currents from transiently transfected HEK293 cells. The pipette solution contained (in mM) 140 NaCl, 10 EGTA, and 10 HEPES, adjusted to pH 7.0 with 1M Tris (osmolarity was ∼296 mmol/kg). The standard extracellular solution contained (in mM): 140 NaCl, 5.4 KCl, 2 CaCl_2_, 0.5 MgCl_2_, 10 D-Glucose, 10 HEPES, and was adjusted to pH 7.3 with 1 M Tris (osmolarity was ∼290 mmol/kg). The divalent-free extracellular solution contained (in mM) 140 NaCl, 10 EDTA, 10 D-Glucose and 10 HEPES, adjusted to pH 7.3 with 1 M Tris (osmolarity was ∼303 mmol/kg). The extracellular solution with Mg^2+^ contained (in mM) 140 NaCl, 5 MgCl_2_ and 10 HEPES, adjusted to pH 7.3 with 1 M Tris (osmolarity was ∼282 mmol/kg). Max Chelator (http://maxchelator.stanford.edu/) was used to calculate concentrations of Mg^2+^, Mg-ATP^2−^, and ATP^4−^ in solutions with 5 mM of MgCl_2_ and 10 µM ATP. Stocks of 10 mM ATP and 1 mM α,β-Me-ATP were prepared in 140 mM NaCl, 10 mM HEPES, pH adjusted to 7.0 with 1 M Tris and were stored as 50-100 µL aliquots to avoid multiple freeze-thaw cycles and working solutions were prepared fresh each day. Concentration-response relations for the WT and mutant receptors were determined by applying 6 different concentrations of ATP (1, 3, 10, 30, 100, and 300 μM) in the standard extracellular solution or in the divalent-free extracellular solution. Membrane currents were recorded under voltage clamp using an Axopatch 200B patch-clamp amplifier (Axon Instruments, Inc.) and digitized online using a Digidata 1550A interface board and pCLAMP 10.5 software (Axon Instruments, Inc.). Currents were filtered at 2 kHz using an eight-pole Bessel filter and digitized at 10 kHz. Solution exchange was achieved using the Rapid Solution Changer RSC-200 (BioLogic), which has the capacity of switching between nine solutions. Pipette resistance was typically 2.5 to 3.5 MΩ; series resistance was less than 6 MΩ and was not compensated.

Concentration–response relations for ATP were obtained for WT and mutant receptor channels, and the Hill equation fit to the data according to the following:

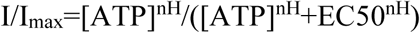

Where I is the normalized current at a given ATP concentration, I_max_ is the maximum normalized current, EC_50_ is the concentration of ATP that evokes half-maximal currents, and nH is the Hill coefficient. Analysis, fitting and plotting of electrophysiological data were done using pClamp 10.5, Origin, Version 2023b (OriginLab Corporation, Northampton, MA, USA) and GraphPad Prism 10. Determination of statistically significant differences (p<0.05) in EC_50_ values was done using Student’s t-test (with and without Welch’s correction).

### Sample Size

Statistical methods were not used to determine sample size. Sample size for cryo-EM studies was determined by availability of microscope time and to ensure sufficient resolution for model building. Sample size for electrophysiological studies was determined empirically by comparing individual measurements with population data obtained under differing conditions until convincing differences or lack thereof were evident. For all electrophysiological experiments, n values represent the number of HEK293 cells studied between 2 to 3 independent experiments.

### Data Exclusions

For electrophysiological experiments, exploratory experiments were undertaken with varying ionic conditions and voltage-clamp protocols to define ideal conditions for measurements reported in this study. Although these preliminary experiments are consistent with the results we report, they were not included in our analysis due to varying experimental conditions (e.g. solution composition and voltage protocols). Once ideal conditions were identified, electrophysiological data were collected for control and mutant constructs until convincing trends in population datasets were obtained. Individual cells were excluded if cells exhibited excessive initial leak currents at the holding voltage (> 200 pA) or if currents arising from expressed channels were too large (>4 nA), resulting in substantial voltage errors or changes in the concentration of ions in the intracellular solutions.

### Randomization and Blinding

Randomization and blinding were not used in this study. The effects of different conditions or mutations on hP2X2Rs heterologously expressed in individual cells was either unambiguously robust or clearly indistinguishable from control conditions.

## Acknowledgments

We thank Tsg-Hui Chang for outstanding assistance and support throughout the duration of this project. We thank, Ana I. Fernández-Mariño, Angela Ballesteros, Shai Silberberg, Mark Mayer, Purushotham Selvakumar, Xiaofeng Tan, Louis Tung Faat Lai and members of the Swartz laboratory for helpful discussion, and Huaibin Wang and Ulrich Baxa in the NIH Multi-Institute Cryo-EM Facility (MICEF) and Zanlin Yu in the NINDS Cryo-EM Core Facility for assistance in acquiring cryo-EM data. This work utilized the NIDDK and NINDS Cryo-EM Core Facilities, the NIH MICEF and computational resources of the NIH HPC Biowulf cluster (http://hpc.nih.gov).

## Author contributions

Conceptualization: SD, MM, KJS

Methodology: SD, MM, KJS

Investigation: SD, MM

Visualization: SD, KJS

Supervision: KJS

Writing—original draft: SD, KJS

Writing—review & editing: SD, MM, KJS

## Data and materials availability

All data are available in the main text or the supplementary materials. Models and/or maps of hP2X2 have been deposited in the Protein Data Bank and/or Electron Microscopy Data Bank (EMDB) with accession codes 9OGH (EMD-70468) for apo hP2X2-L in nanodiscs with PIP2 in C3, 9OIR (EMD-70527) for hP2X2-L in detergent with residual ATP in C3, EMD- 70526 for hP2X2-L in detergent treated with apyrase with residual ATP in C3, EMD-70457 for apo hP2X2-S in nanodiscs in C3, EMD-70456 for hP2X2-S with Mg^2+^ in nanodiscs in C3, 9Z32 (EMD-73782) for hP2X2-S with Mg-ATP^2-^ in nanodiscs in C3, 9OHK (EMD-70496) for hP2X2-L with suramin in nanodiscs with PIP2 in C3. For datasets for hP2X2-L with free-ATP in nanodiscs with Na^+^ and PIP2 in C3, 9ON5/ 9ON6 (EMD-70629) for class 1, 9OJK (EMD-70543) for class 2, 9OM0 (EMD-70602) for class 3 and 9OMR (EMD-70617) for class 4.

