## Supplementary Materials for "Mechanisms of ligand recognition and channel opening for P2X2 receptors in lipid nanodiscs"

**This PDF file includes:**

Figs. S1 to S10
Tables S1 to S3
Movies S1 to S2

**Other Supplementary Materials for this manuscript include the following:**

Movies S1 to S2

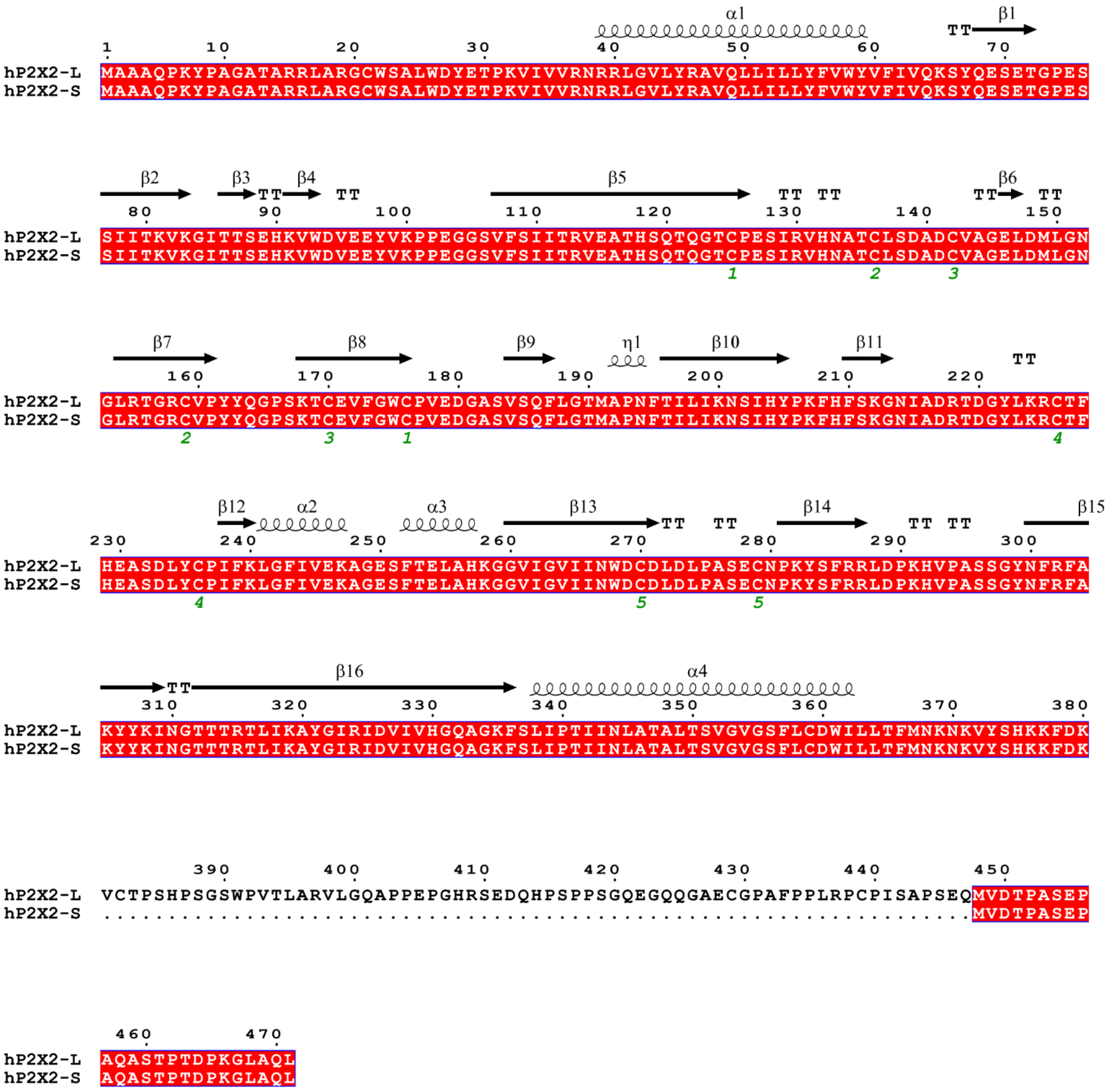

**Fig. S1. Sequence alignment for two splice variants of hP2X2Rs.**
Sequence alignment of hP2X2-L and hP2X2-S receptor channels was performed using Clustal Omega [75]. Secondary structural features are indicated above the sequence are from the apo structure of hP2X2-L and detected using ESPript [76]. TT indicates a turn. Cys involved in disulfide bonds are indicated with green numbers. The two splice variants differ only near the C-terminal end of the protein, between V381-Q447 (67 residues).

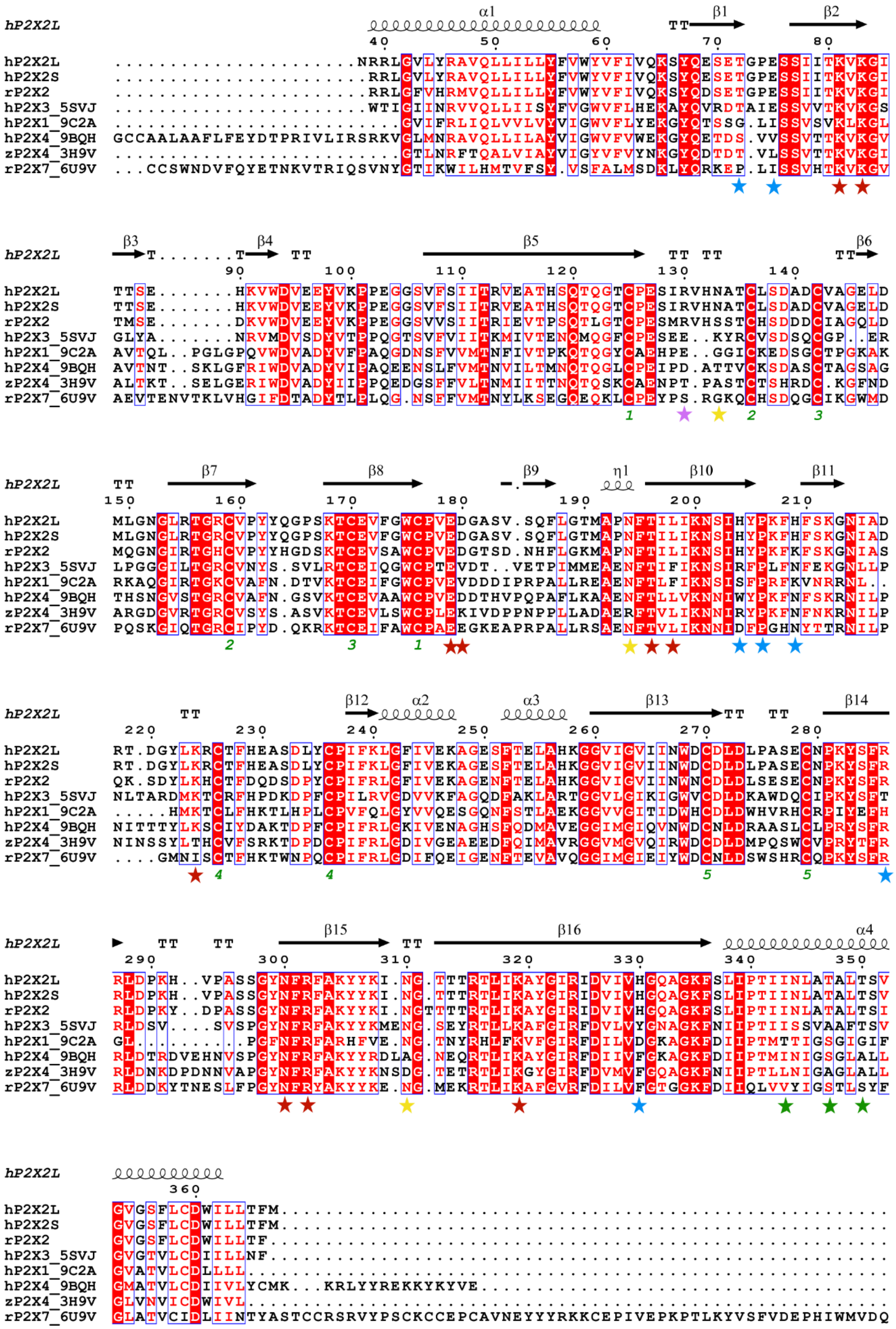
hP2X2L
α1 β1 β2
40 50 60 70 80
hP2X2L ......................NRRLGVLYRAVQLLILLYFVWYVFIVQKSYQESETGPESSIITKVKGI
hP2X2S .......................RRLGVLYRAVQLLILLYFVWYVFIVQKSYQESETGPESSIITKVKGI
rP2X2 .......................RRLGFVHRMVVQLLILLYFVWYVFIVQKSYQDSETGPESSIITKVKGI
hP2X3_5SVJ .......................WTIGIINRVVQLLIISYFVGWVFLHEKAYQVRDTAIESSVVTKVKGS
hP2X1_9C2A .........................GVIFRLIQLVVLVYVIGWVFLYEKGYQTSSG.LISSVSVKLKGL
hP2X4_9BQH GCCAALAAFLFEYDTPRIVLIRSRKVGLMNRAVQLLILAYVIGWVFVWEKGYQETDS.VVSSVTTKVKGV
zP2X4_3H9V .........................GTLNRFTQALVIAYVIGYVFVYNKGYQDTDT.VLSSVTTKVKGI
rP2X7_6U9V ...CCSWNDVFQYETNKVTRIQSVNYGTIKWILHMTVFSY.VSFALMSDKLYQRKEP.LISSVHTKVKGV
hP2X2L
β3 β4 β5 β6
90 100 110 120 130 140
hP2X2L TTSE.......HKVWDVEEYVKPPEGGSVFSIITRVEATHSQTQGTCPESIRVHNATCLSDADCVAGELD
hP2X2S TTSE.......HKVWDVEEYVKPPEGGSVFSIITRVEATHSQTQGTCPESIRVHNATCLSDADCVAGELD
rP2X2 TMSE.......DKVWDVEEYVKPPEGGSVVSIITRIEVTPSQTLGTCPESMRVHSSTCHSDDDCIAGQLD
hP2X3_5SVJ GLYA.......NRVMDVSDYVTPPQGTSVFVIITKMIVTENQMQGFCPESEE..KYRCVSDSQCGP..ER
hP2X1_9C2A AVTQL..PGLGPQVWDVADYVFPAQGDNSFVVMTNFIVTPKQTQGYCAEHPE..GGICKEDSGCTPGKAK
hP2X4_9BQH AVTNT..SKLGFRIWDVADYVIPAQEENSLFVMTNVILTMNQTQGLCPEIPD.ATTVCKSDASCTAGSAG
zP2X4_3H9V ALTKT..SELGERIWDVADYIIPPQEDGSFFVLTNMIITTNQTQSKCAENPT.PASTCTSHRDC.KGFND
rP2X7_6U9V AEVTENVTKLVHGIFDTADYTLPLQG.NSFFVMTNYLKSEGQEQKLCPEYPS.RGKQCHSDQGCIKGWMD
hP2X2L
β7 β8 β9 η1 β10 β11
150 160 170 180 190 200 210
hP2X2L MLGNGLRTGRCVPYYQGPSKTCEVFGWCPVEDGASV.SQFLGTMAPNFTILIKNSIHYPKFHFSKGNIAD
hP2X2S MLGNGLRTGRCVPYYQGPSKTCEVFGWCPVEDGASV.SQFLGTMAPNFTILIKNSIHYPKFHFSKGNIAD
rP2X2 MQGNGIRTGHCVPYYHGDSKTCEVSAWCPVEDGTSD.NHFLGKMAPNFTILIKNSIHYPKFKFSKGNIAS
hP2X3_5SVJ LPGGGILTGRCVNYS.SVLRTCEIQGWCPTEVDT..VETPIMMEAENFTIFIKNSIRFPLFNFEKGNLLP
hP2X1_9C2A RKAQGIRTGKCVAFN.DTVKTCEIFGWCPVEVDDDIPRPALLREAENFTLFIKNSISFPRFKVNRRNL..
hP2X4_9BQH THSNGVSTGRCVAFN.GSVKTCEVAAWCPVEDDTHVPQPAFLKAAENFTLLVKNNIWYPKFNFSKRNILP
zP2X4_3H9V ARGDGVRTGRCVSYS.ASVKTCEVLSWCPLEKIVDPPNPPLLADAERFTVLIKNNIRYPKFNFNKRNILP
rP2X7_6U9V PQSKGIQTGRCIPYD.QKRKTCEIFAWCPAEEGKEAPRPALLRSAENFTVLIKNNIDFPGHNYTTRNILP
hP2X2L
β12 α2 α3 β13 β14
220 230 240 250 260 270 280
hP2X2L RT.DGYLKRCTFHEASDLYCPIFKLGFIVEKAGESFTELAHKGGVIGVIINWDCDLDLPASECNPKYSFR
hP2X2S RT.DGYLKRCTFHEASDLYCPIFKLGFIVEKAGESFTELAHKGGVIGVIINWDCDLDLPASECNPKYSFR
rP2X2 QK.SDYLKHCTFDQDSDPYCPIFRLGFIVEKAGENFTELAHKGGVIGVIINWNCDLDLSESECNPKYSFR
hP2X3_5SVJ NLTARDMKTCRFHPDKDPFCPILRVGDVVKFAGQDFAKLARTGGVLGIKIGWVCDLDKAWDQCIPKYSFT
hP2X1_9C2A .....HMKTCLFHKTLHPLCPVFQLGYVVQESGQNFSTLAEKGGVVGITIDWHCDLDWHVRHCRPIYEFH
hP2X4_9BQH NITTTYLKSCIYDAKTDPFCPIFRLGKIVENAGHSFQDMAVEGGIMGIQVNWDCNLDRAASLCLPRYSFR
zP2X4_3H9V NINSSYLTHCVFSRKTDPDCPIFRLGDIVGEAEEDFQIMAVRGGVMGVQIRWDCDLDMPQSWCVPRYTFR
rP2X7_6U9V ....GMNISCTFHKTWNPQCPIFRLGDIFQEIGENFTEVAVQGGIMGIEIYWDCNLDSWSHRCQPKYSFR
hP2X2L
β15 β16 α4
290 300 310 320 330 340 350
hP2X2L RLDPKH..VPASSGYNFRFAKYYKI.NG.TTTRTLIKAYGIRIDVIVHGQAGKFSLIPTIINLATALTSV
hP2X2S RLDPKH..VPASSGYNFRFAKYYKI.NG.TTTRTLIKAYGIRIDVIVHGQAGKFSLIPTIINLATALTSV
rP2X2 RLDPKY..DPASSGYNFRFAKYYKI.NGTTTTRTLIKAYGIRIDVIVHGQAGKFSLIPTIINLATALTSI
hP2X3_5SVJ RLDSV....SVSPGYNFRFAKYYKMENG.SEYRTLLKAFGIRFDVLVYGNAGKFNIIPTIISSVAAFTSV
hP2X1_9C2A GL.........PGFNFRFARHFVE.NG.TNYRHLFKVFGIRFDILVDGKAGKFDIIPTMTTIGSGIGIF
hP2X4_9BQH RLDTRDVEHNVSPGYNFRFAKYYRDLAG.NEQRTLIKAYGIRFDIIVFGKAGKFDIIPTMINIGSGLALL
zP2X4_3H9V RLDNKDPDNNVAPGYNFRFAKYYKNSDG.TETRTLIKGYGIRFDVMVFGQAGKFNIIPTLLNIGAGLALL
rP2X7_6U9V RLDDKYTNESLFPGYNFRYAKYYKE.NG.MEKRTLIKAFGVRFDILVFGTGGKFDIIQLVVYIGSTLSYF
hP2X2L
360
hP2X2L GVGSFLCDWILLTFM.......................................................
hP2X2S GVGSFLCDWILLTFM.......................................................
rP2X2 GVGSFLCDWILLTF........................................................
hP2X3_5SVJ GVGTVLCDIILLNF........................................................
hP2X1_9C2A GVATVLCDLLLL..........................................................
hP2X4_9BQH GMATVLCDIIVLYCMK...KRLYYREKKYKYVE.....................................
zP2X4_3H9V GLVNVICDWIVL..........................................................
rP2X7_6U9V GLATVCIDLIINTYASTCCRSRVYPSCKCCEPCAVNEYYYRKKCEPIVEPKPTLKYVSFVDEPHIWMVDQ

**🡐 Fig. S2. Structure-based sequence alignment for P2XRs in apo state.**

Structure-based sequence alignment of P2XRs (hP2X2-L, hP2X2-S and rP2X2-L are from this study, and for all others PDB IDs are provided in the figure). Secondary structural features are indicated above the sequence are from the apo structure of hP2X2-L and detected using ESPript [76]. TT indicates a turn. Residues involved in ATP binding are indicated with a red star. R130 in P2X2 is indicated with a pink star. Residues involved in coupling ATP binding to channel opening are indicated with a blue star. Glycosylated Asn residues are indicated with a yellow star. Residues in the activation gate are indicated with a green star. Cys involved in disulfide bonds are indicated with green numbers.

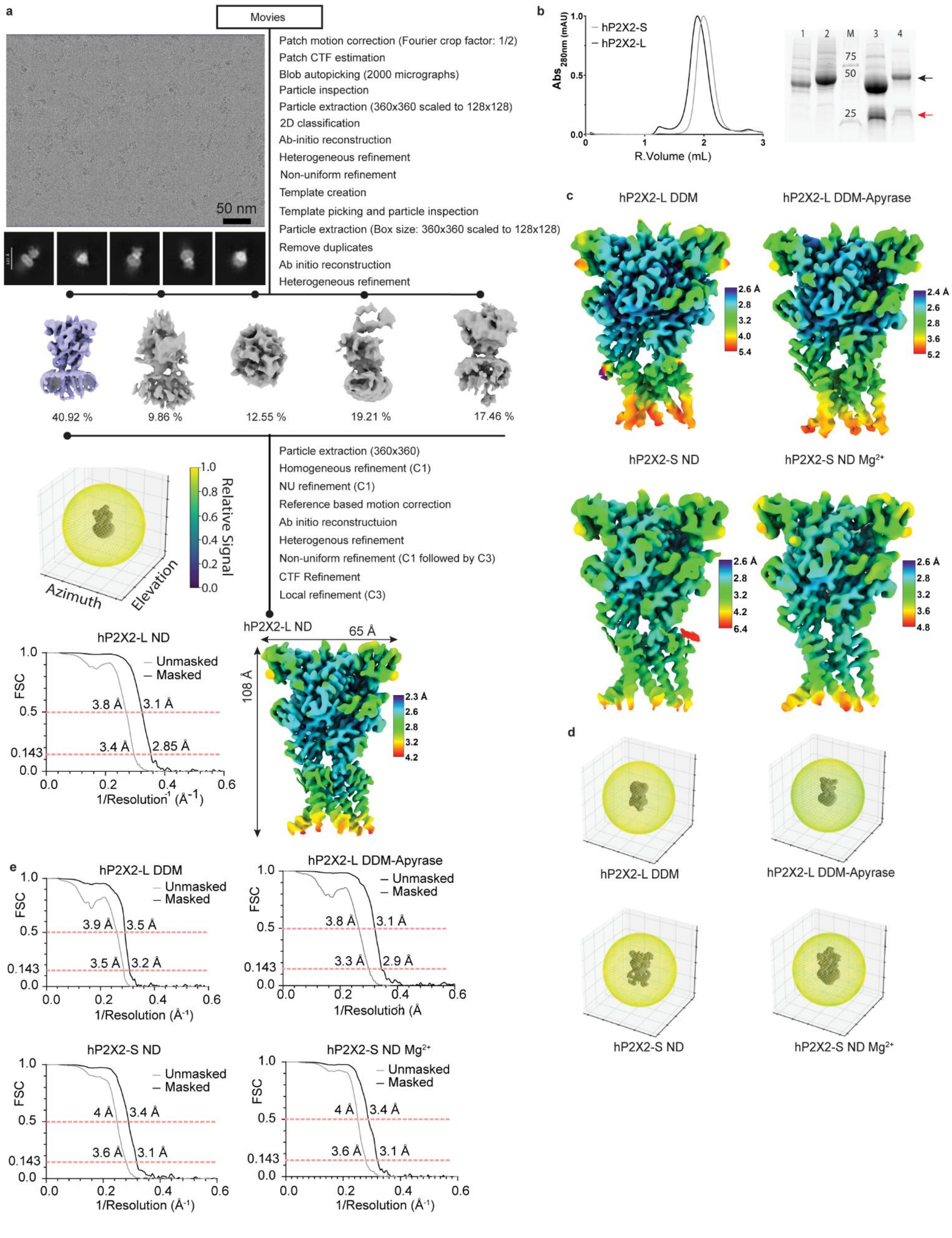
a
Movies
Patch motion correction (Fourier crop factor: 1/2)
Patch CTF estimation
Blob autopicking (2000 micrographs)
Particle inspection
Particle extraction (360x360 scaled to 128x128)
2D classification
Ab-initio reconstruction
Heterogeneous refinement
Non-uniform refinement
Template creation
Template picking and particle inspection
Particle extraction (Box size: 360x360 scaled to 128x128)
Remove duplicates
Ab initio reconstruction
Heterogeneous refinement
50 nm
40.92 %
9.86 %
12.55 %
19.21 %
17.46 %
Particle extraction (360x360)
Homogeneous refinement (C1)
NU refinement (C1)
Reference based motion correction
Ab initio reconstructuion
Heterogenous refinement
Non-uniform refinement (C1 followed by C3)
CTF Refinement
Local refinement (C3)
Relative Signal
Azimuth
Elevation
hP2X2-L ND
65 Å
108 Å
hP2X2-L ND
Unmasked
Masked
FSC
3.8 Å
3.1 Å
3.4 Å
2.85 Å
1/Resolution (Å-1)
b
hP2X2-S
hP2X2-L
Abs 280nm (mAU)
R.Volume (mL)
1 2 M 3 4
75
50
25
c
hP2X2-L DDM
hP2X2-L DDM-Apyrase
hP2X2-S ND
hP2X2-S ND Mg2+
d
hP2X2-L DDM
hP2X2-L DDM-Apyrase
hP2X2-S ND
hP2X2-S ND Mg2+
e
hP2X2-L DDM
3.9 Å
3.5 Å
3.5 Å
3.2 Å
hP2X2-L DDM-Apyrase
3.8 Å
3.1 Å
3.3 Å
2.9 Å
hP2X2-S ND
4 Å
3.4 Å
3.6 Å
3.1 Å
hP2X2-S ND Mg2+
4 Å
3.4 Å
3.6 Å
3.1 Å

**← Fig. S3. Cryo-EM imaging for apo hP2X2.**

**a**) Data processing workflow for the cryo-EM structures of apo hP2X2. Illustrative micrograph and 2D class averages are shown to the left. Local resolution unsharpened map for the structure of hP2X2-L in nanodiscs (ND) is shown alongside direction distribution plots of the 3D reconstruction illustrating the distribution of particles in different orientations and Fourier Shell Correlation (FSC) curves: FSC calculated without mask (grey) and FSC calculated using the tight mask with correction by noise substitution (black). **b**) Superose 6 Increase (10/150 GL) size-exclusion chromatograms for purified samples of hP2X2-S and hP2X2-L reconstituted into nanodiscs used for preparing grids. SDS-PAGE stain-free gel (Biorad) to the right where samples were: 1) hP2X2-S in DDM, 2) hP2X2-L in DDM, 3) hP2X2-S in nanodiscs and 4) hP2X2-L in nanodiscs with $Na^+$ as the primary cation (as was used in all other samples). Black arrow indicates hP2X2R (hP2X2-S: ~44.8kDa, hP2X2-L: ~51.8 kDa) and red arrow indicates MSP1E3D1 (~32.6 kDa). **c**) Local resolution unsharpened maps for four different structures solved without adding ATP using a FSC of 0.5. **d)** Direction distribution plots of the 3D reconstructions illustrating the distribution of particles in different orientations for the same structures shown in panel c. **e)** Fourier Shell Correlation (FSC) curves for the same structures shown in panel c: FSC calculated without mask (grey) and FSC calculated using the tight mask with correction by noise substitution (black).

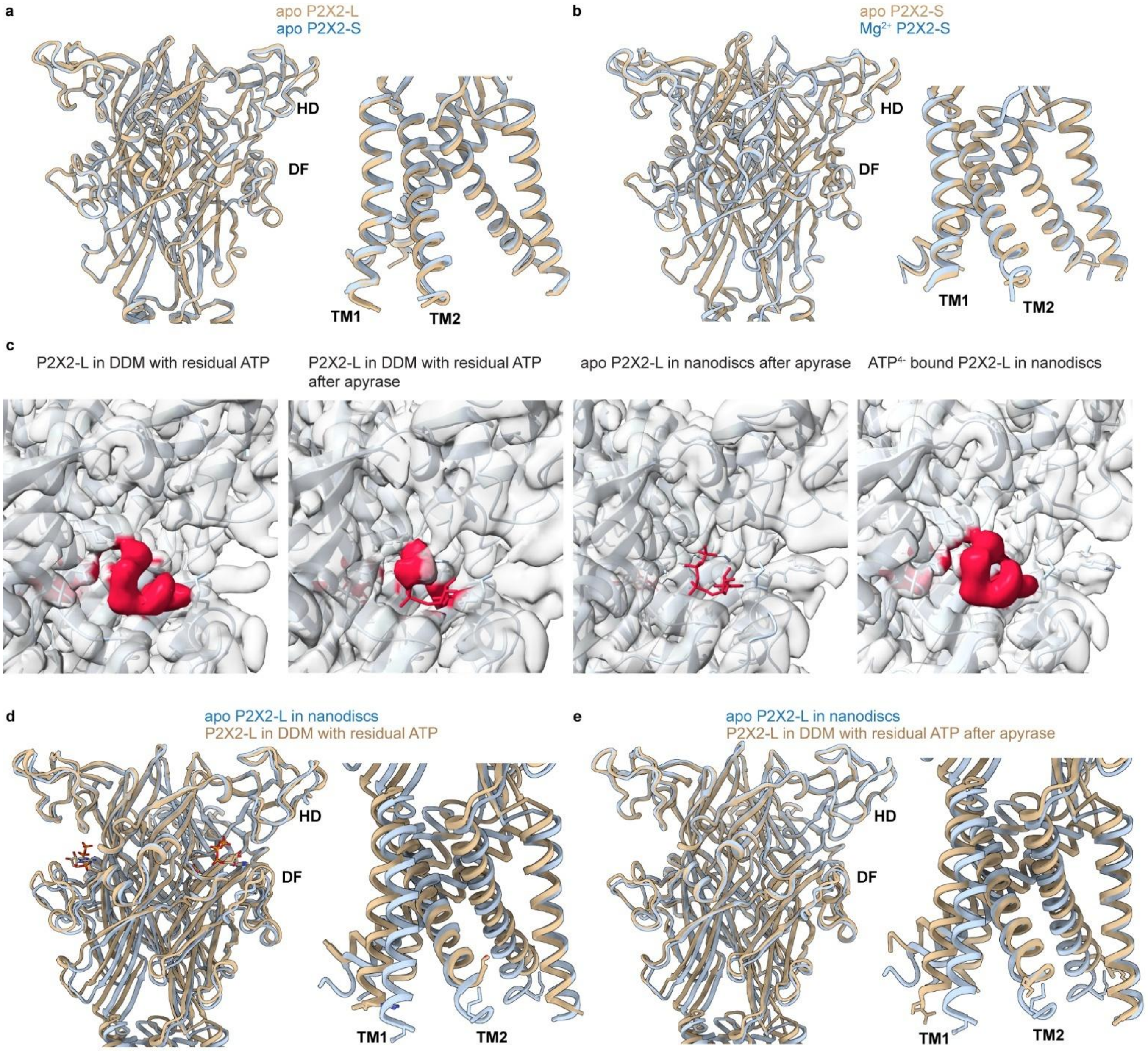


**Fig. S4. Comparison of structures of hP2X2 solved without ATP added to the samples.**
**a**) Superimposed structures in the extracellular and TMD for apo hP2X2-L (tan) with apo hP2X2-S (blue). **b**) Superimposed structures in the extracellular and TMD for apo hP2X2-S (tan) with hP2X2-S solved in the presence of 5 mM $Mg^{2+}$ (blue). **c**) Cryo-EM unsharpened maps for structures of hP2X2-L solved in DDM without or with apyrase treatment, reconstituted into nanodiscs with apyrase treatment or with $ATP^{4-}$. Density assigned to ATP is shown in red and for apo the position where $ATP^{4-}$ would be bound to hP2X2-L is indicated by red stick representation. **d**) Superimposed structures in the extracellular and TMD for apo hP2X2-L in nanodiscs (blue) with hP2X2-L in DDM with residual ATP remaining bound during purification (tan). **e**) Superimposed structures in the extracellular and TMD for apo hP2X2-L in nanodiscs (blue) with hP2X2-L in DDM with residual ATP remaining bound during purification even after treatment with apyrase (tan).

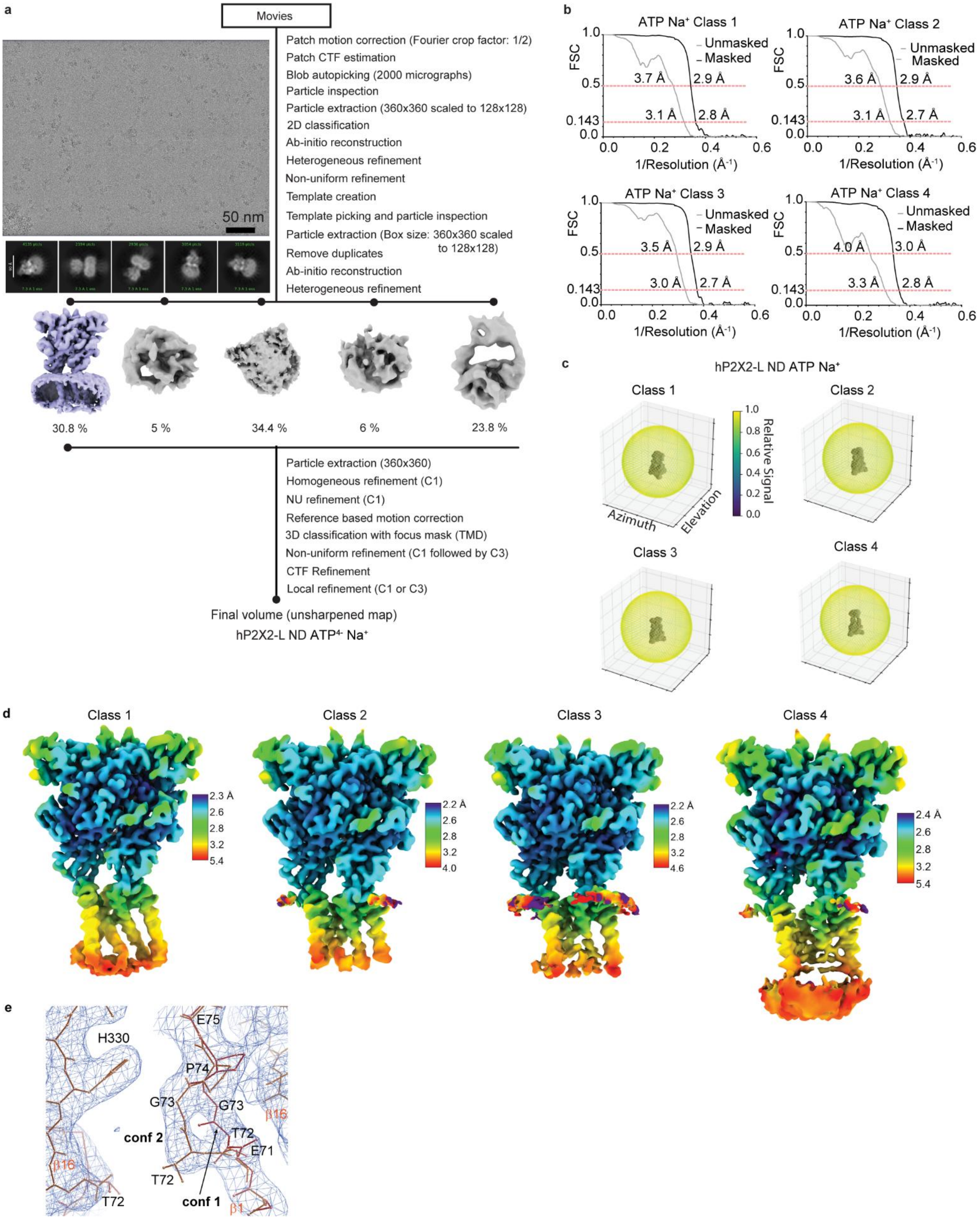
a
Movies
Patch motion correction (Fourier crop factor: 1/2)
Patch CTF estimation
Blob autopicking (2000 micrographs)
Particle inspection
Particle extraction (360x360 scaled to 128x128)
2D classification
Ab-initio reconstruction
Heterogeneous refinement
Non-uniform refinement
Template creation
Template picking and particle inspection
Particle extraction (Box size: 360x360 scaled to 128x128)
Remove duplicates
Ab-initio reconstruction
Heterogeneous refinement
50 nm
30.8 %
5 %
34.4 %
6 %
23.8 %
Particle extraction (360x360)
Homogeneous refinement (C1)
NU refinement (C1)
Reference based motion correction
3D classification with focus mask (TMD)
Non-uniform refinement (C1 followed by C3)
CTF Refinement
Local refinement (C1 or C3)
Final volume (unsharpened map)
hP2X2-L ND ATP4- Na+
b
ATP Na+ Class 1
ATP Na+ Class 2
ATP Na+ Class 3
ATP Na+ Class 4
FSC
Unmasked
Masked
1/Resolution (Å-1)
3.7 Å
2.9 Å
3.1 Å
2.8 Å
3.6 Å
2.9 Å
3.1 Å
2.7 Å
3.5 Å
2.9 Å
3.0 Å
2.7 Å
4.0 Å
3.0 Å
3.3 Å
2.8 Å
c
hP2X2-L ND ATP Na+
Class 1
Class 2
Class 3
Class 4
Relative Signal
Azimuth
Elevation
d
Class 1
Class 2
Class 3
Class 4
e
H330
E75
P74
G73
G73
T72
E71
conf 2
conf 1
T72
T72
β16
β16
β1

**🡐 Fig. S5. Cryo-EM imaging for hP2X2 with ATP$^{4-}$.**

**a**) Data processing workflow for the cryo-EM structures of hP2X2 with ATP$^{4-}$. Illustrative micrograph and 2D class averages are shown to the left. Local resolution maps for the structures of 100 μM ATP$^{4-}$ (plus 5 mM EDTA) bound hP2X2-L in nanodiscs (ND) for the four different classes solved in $Na^+$. **b)** Fourier Shell Correlation (FSC) curves for the same structures shown in panel d: FSC calculated without mask (grey) and FSC calculated using the tight mask with correction by noise substitution (black). **c)** Direction distribution plots of the 3D reconstructions illustrating the distribution of particles in different orientations for the same structures shown in panel d. **d**) Local resolution unsharpened maps for the four classes determined with ATP$^{4-}$ using a FSC of 0.5. **e**) Cryo-EM map of the β1/β16 interface showing two conformations of the T72-P74 segment of β1. For conformation 2, the β1 strand is intact, as observed in other structures of P2XRs, whereas in conformation 1, T72 and G73 are located closer to the subunit interface where T72 interacts with the same position in the apo state.

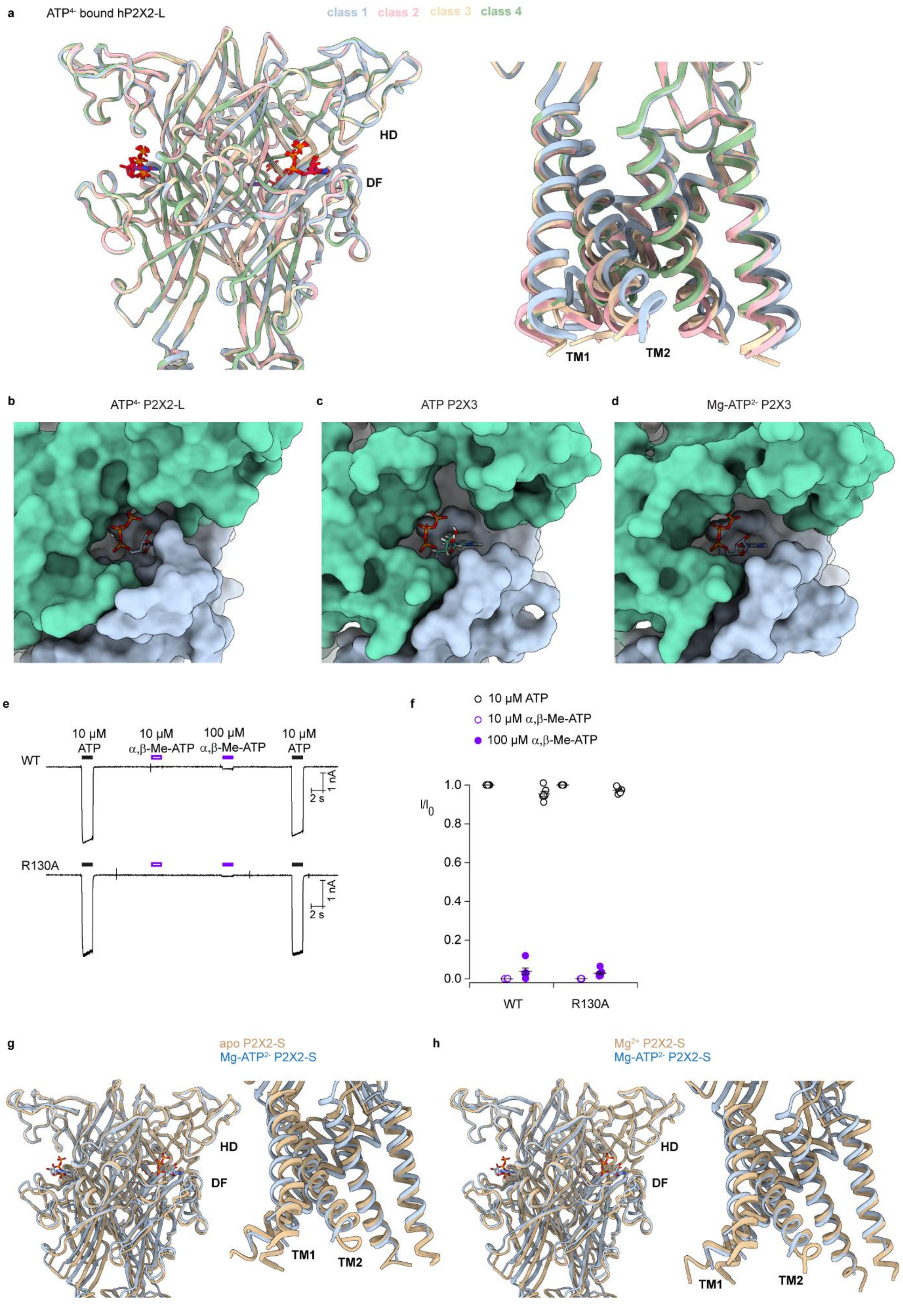
a
ATP4- bound hP2X2-L
class 1 class 2 class 3 class 4
HD
DF
TM1
TM2
b
ATP4- P2X2-L
c
ATP P2X3
d
Mg-ATP2- P2X3
e
10 μM ATP
10 μM α,β-Me-ATP
100 μM α,β-Me-ATP
10 μM ATP
WT
R130A
1 nA
2 s
f
10 μM ATP
10 μM α,β-Me-ATP
100 μM α,β-Me-ATP
I/I0
1.0
0.8
0.6
0.4
0.2
0.0
WT
R130A
g
apo P2X2-S
Mg-ATP2- P2X2-S
HD
DF
TM1
TM2
h
Mg2+ P2X2-S
Mg-ATP2- P2X2-S
HD
DF
TM1
TM2

**← Fig. S6. Comparison of structures of hP2X2 solved in the presence of varying ligands and ionic conditions.**

**a**) Superimposed structures in the ECD and TMD for the four classes resolved with $ATP^{4-}$ bound to hP2X2-L in nanodiscs **b-d**) Surface renderings of the ATP binding site with $ATP^{4-}$ bound to hP2X2-L (class 2), ATP bound to hP2X3 (5svk) and Mg-$ATP^{2-}$ bound to hP2X3 (6ah4), respectively. **e**) Currents elicited at a holding voltage of −60 mV for cells expressing WT hP2X2-L and R130A applying external solutions containing 10 μM ATP before and after applying either 10 μM or 100 μM α,β-Me-ATP in a physiological solution containing 2 mM $Ca^{2+}$ and 0.5 mM $Mg^{2+}$. **f**) Plots of normalized currents activated by ATP or α,β-Me-ATP for WT and R130A. Individual cells are shown as open symbols with mean and S.E.M. shown as bars with n=6 in 2 independent experiments for WT and R130A. **g**) Superimposed structures in the ECD and TMD for apo hP2X2-S (tan) with Mg-$ATP^{2-}$ bound hP2X2-S (blue). **h**) Superimposed structures of the ECD and TMD for hP2X2-S solved in the presence of 5 mM $Mg^{2+}$ (tan) with Mg-$ATP^{2-}$bound hP2X2-S (blue).

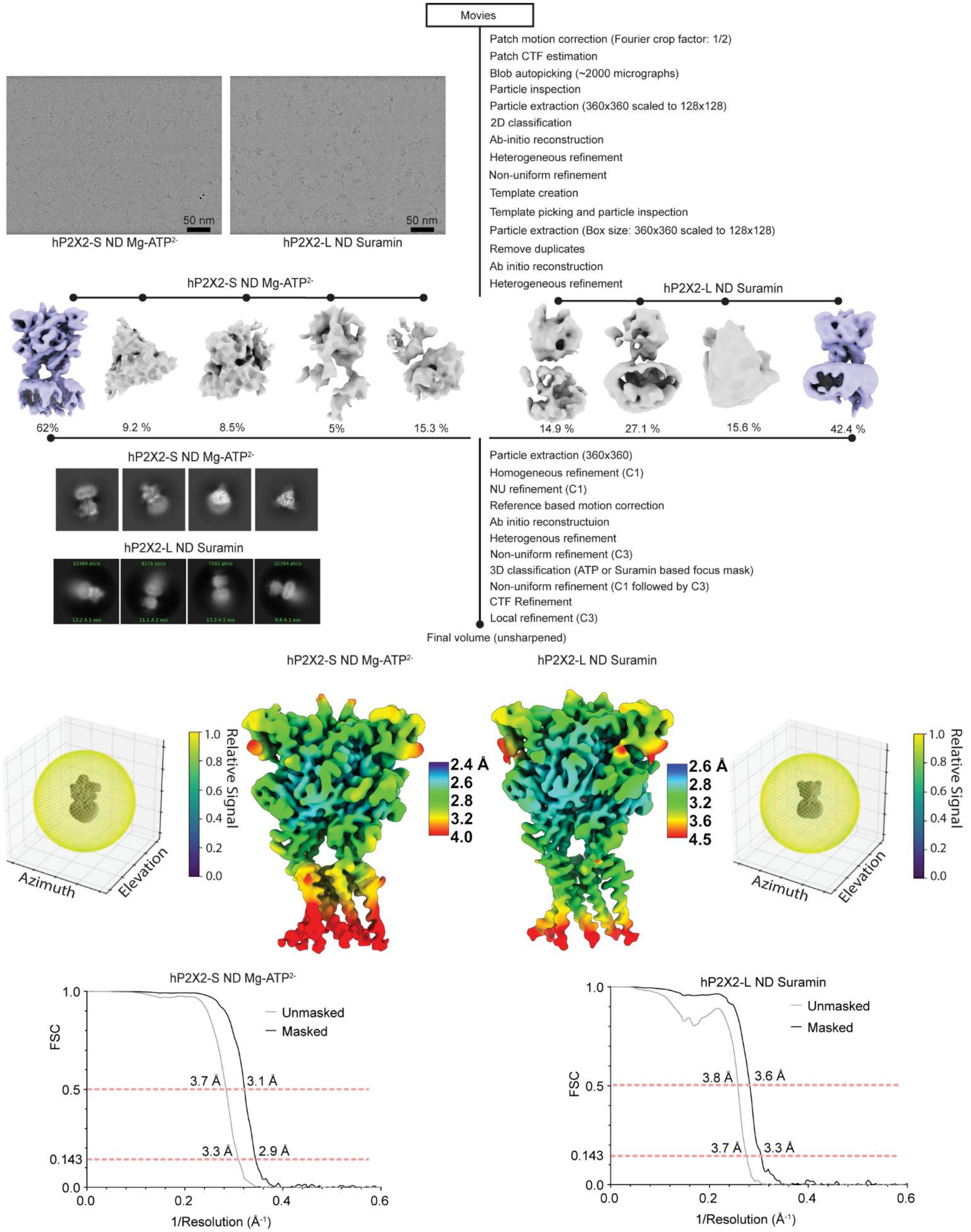

Movies
Patch motion correction (Fourier crop factor: 1/2)
Patch CTF estimation
Blob autopicking (~2000 micrographs)
Particle inspection
Particle extraction (360x360 scaled to 128x128)
2D classification
Ab-initio reconstruction
Heterogeneous refinement
Non-uniform refinement
Template creation
Template picking and particle inspection
Particle extraction (Box size: 360x360 scaled to 128x128)
Remove duplicates
Ab initio reconstruction
Heterogeneous refinement
50 nm
50 nm
hP2X2-S ND Mg-ATP2-
hP2X2-L ND Suramin
hP2X2-S ND Mg-ATP2-
hP2X2-L ND Suramin
62%
9.2 %
8.5%
5%
15.3 %
14.9 %
27.1 %
15.6 %
42.4 %
hP2X2-S ND Mg-ATP2-
hP2X2-L ND Suramin
Particle extraction (360x360)
Homogeneous refinement (C1)
NU refinement (C1)
Reference based motion correction
Ab initio reconstructuion
Heterogenous refinement
Non-uniform refinement (C3)
3D classification (ATP or Suramin based focus mask)
Non-uniform refinement (C1 followed by C3)
CTF Refinement
Local refinement (C3)
Final volume (unsharpened)
hP2X2-S ND Mg-ATP2-
hP2X2-L ND Suramin
Azimuth
Elevation
Relative Signal
1.0
0.8
0.6
0.4
0.2
0.0
2.4 Å
2.6
2.8
3.2
4.0
2.6 Å
2.8
3.2
3.6
4.5
Azimuth
Elevation
Relative Signal
1.0
0.8
0.6
0.4
0.2
0.0
hP2X2-S ND Mg-ATP2-
FSC
Unmasked
Masked
3.7 Å
3.1 Å
3.3 Å
2.9 Å
1.0
0.5
0.143
0.0
0.0
0.2
0.4
0.6
1/Resolution (Å-1)
hP2X2-L ND Suramin
FSC
Unmasked
Masked
3.8 Å
3.6 Å
3.7 Å
3.3 Å
1.0
0.5
0.143
0.0
0.0
0.2
0.4
0.6
1/Resolution (Å-1)

**← Fig. S7. Cryo-EM imaging for hP2X2 with Mg-ATP$^{2-}$ or suramin bound.**

Data processing workflow for the cryo-EM structures of hP2X2-S with Mg-ATP$^{2-}$ bound and hP2X2-L with suramin bound. Illustrative micrographs and 2D class averages are shown to the left. Local resolution unsharpened maps (FSC of 0.5) are shown alongside direction distribution plots of the 3D reconstruction illustrating the distribution of particles in different orientations. Fourier Shell Correlation (FSC) curves for the same structures are shown: FSC calculated without mask (grey) and FSC calculated using the tight mask with correction by noise substitution (black).

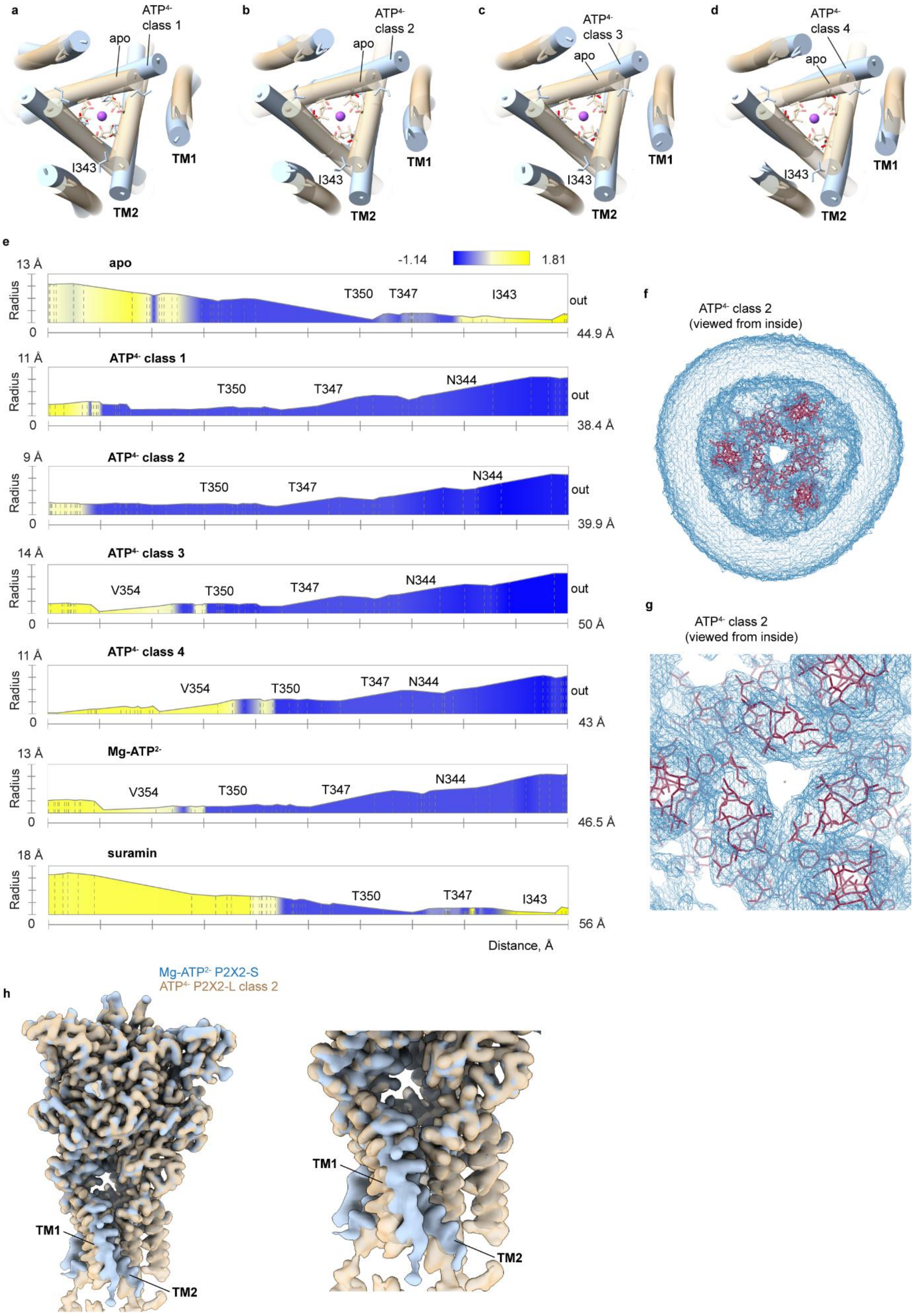
a
ATP4- class 1
apo
I343
TM1
TM2
b
ATP4- class 2
c
ATP4- class 3
d
ATP4- class 4
e
apo
-1.14
1.81
Radius
T350
T347
I343
out
44.9 Å
ATP4- class 1
N344
38.4 Å
ATP4- class 2
39.9 Å
ATP4- class 3
V354
50 Å
ATP4- class 4
43 Å
Mg-ATP2-
46.5 Å
suramin
56 Å
Distance, Å
f
ATP4- class 2
(viewed from inside)
g
ATP4- class 2
(viewed from inside)
h
Mg-ATP2- P2X2-S
ATP4- P2X2-L class 2
TM1
TM2

**← Fig. S8. Structures of hP2X2 with $ATP^{4-}$ bound.**

**a-d**) Superimposition of the structures in the TM regions for apo (tan) with a $Na^+$ ion (purple sphere) and class 1 to 4 of $ATP^{4-}$ bound (blue) for hP2X2-L viewed from the extracellular side of the membrane. Residues in the gate region are shown as stick representation. **e**) Radius of the pore plotted against distance along the pore perpendicular to the plane of the membrane with intracellular at 0 Å obtained using MOLE [73]. Color coding is for hydrophobicity of pore lining residues. **f**) Cryo-EM density for $ATP^{4-}$ bound hP2X2-L (class 2) in lipid nanodiscs viewed from an intracellular perspective. **g**) Zoomed in view of the pore from the same perspective as in f. **h)** Superimposed cryo-EM maps (DeepEMhancer sharpened) for $ATP^{4-}$ (class 2, tan) and Mg-$ATP^{2-}$ (blue) bound hP2X2 in lipid nanodiscs showing distinct movements of the TM1 and TM2 helices.

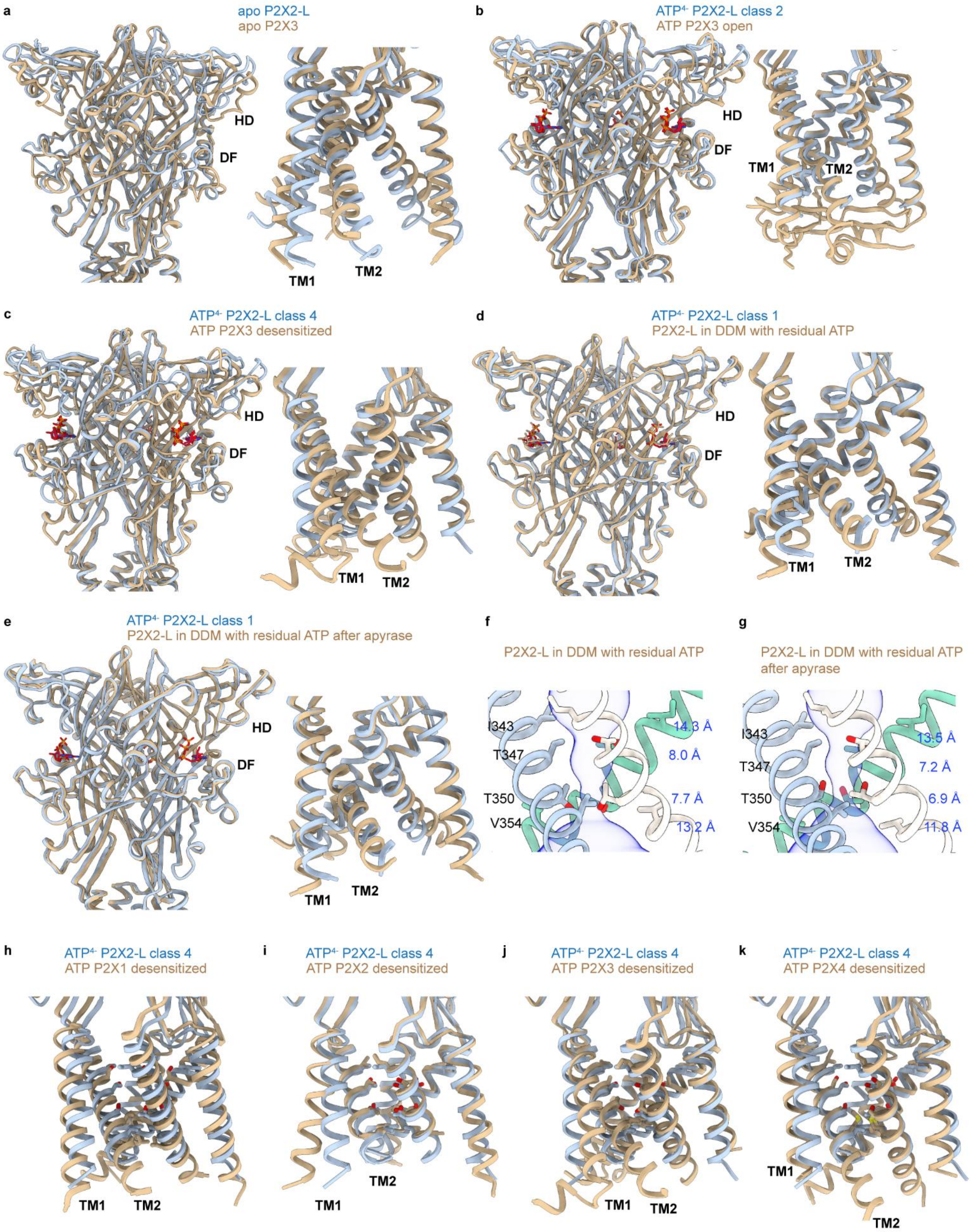
a
apo P2X2-L
apo P2X3
HD
DF
TM1
TM2
b
ATP4- P2X2-L class 2
ATP P2X3 open
HD
DF
TM1
TM2
c
ATP4- P2X2-L class 4
ATP P2X3 desensitized
HD
DF
TM1
TM2
d
ATP4- P2X2-L class 1
P2X2-L in DDM with residual ATP
HD
DF
TM1
TM2
e
ATP4- P2X2-L class 1
P2X2-L in DDM with residual ATP after apyrase
HD
DF
TM1
TM2
f
P2X2-L in DDM with residual ATP
I343
T347
T350
V354
14.3 Å
8.0 Å
7.7 Å
13.2 Å
g
P2X2-L in DDM with residual ATP after apyrase
I343
T347
T350
V354
13.5 Å
7.2 Å
6.9 Å
11.8 Å
h
ATP4- P2X2-L class 4
ATP P2X1 desensitized
TM1
TM2
i
ATP4- P2X2-L class 4
ATP P2X2 desensitized
TM1
TM2
j
ATP4- P2X2-L class 4
ATP P2X3 desensitized
TM1
TM2
k
ATP4- P2X2-L class 4
ATP P2X4 desensitized
TM1
TM2

**← Fig. S9. Comparison of structures of hP2X2 in nanodiscs with hP2XRs in detergent.**

**a)** Superimposed structures of apo hP2X2-L in lipid nanodiscs with apo hP2X3 in detergent (5svj). Differences are largely restricted to the head and L-flipper regions with the ECD and within the TMD. **b)** Superimposed structures of $ATP^{4-}$ bound hP2X2-L (class 2; open) in lipid nanodiscs with ATP bound open hP2X3 in detergent (5svk). Differences are largely restricted to the head and L-flipper regions with the ECD and within the TMD. **c)** Superimposed structures of $ATP^{4-}$ bound hP2X2-L (class 4; desensitized) in lipid nanodiscs with ATP bound desensitized hP2X3 in detergent (5svl). Differences are largely restricted to the head and L-flipper regions with the ECD and within the TMD. **d**) Superimposed structures of $ATP^{4-}$ bound hP2X2-L in nanodiscs (class 1) with hP2X2-L in DDM. **e**) Superimposed structures of $ATP^{4-}$ bound hP2X2-L in nanodiscs (class 1) with hP2X2-L in DDM after apyrase treatment. **f,g**) MOLE representations for the pore of hP2X2-L in DDM without and with apyrase treatment. Residues in the gate region are shown as stick representation with distances shown between adjacent C$\alpha$ atoms. **h)** Superimposed structures of $ATP^{4-}$ bound hP2X2-L (class 4; desensitized) in lipid nanodiscs with ATP bound desensitized hP2X1 in detergent (9c2b). **i)** Superimposed structures of $ATP^{4-}$ bound hP2X2-L (class 4; desensitized) in lipid nanodiscs with ATP bound desensitized hP2X2 in detergent (9ddw). **j)** Superimposed structures of $ATP^{4-}$ bound hP2X2-L (class 4; desensitized) in lipid nanodiscs with ATP bound desensitized hP2X3 in detergent (5svl). **k)** Superimposed structures of $ATP^{4-}$ bound hP2X2-L (class 4; desensitized) in lipid nanodiscs with ATP bound desensitized hP2X4 in detergent (9c48). For panels h-k, residues equivalent to I343, T347, T350 and V354 in hP2X2 are shown in stick representation.

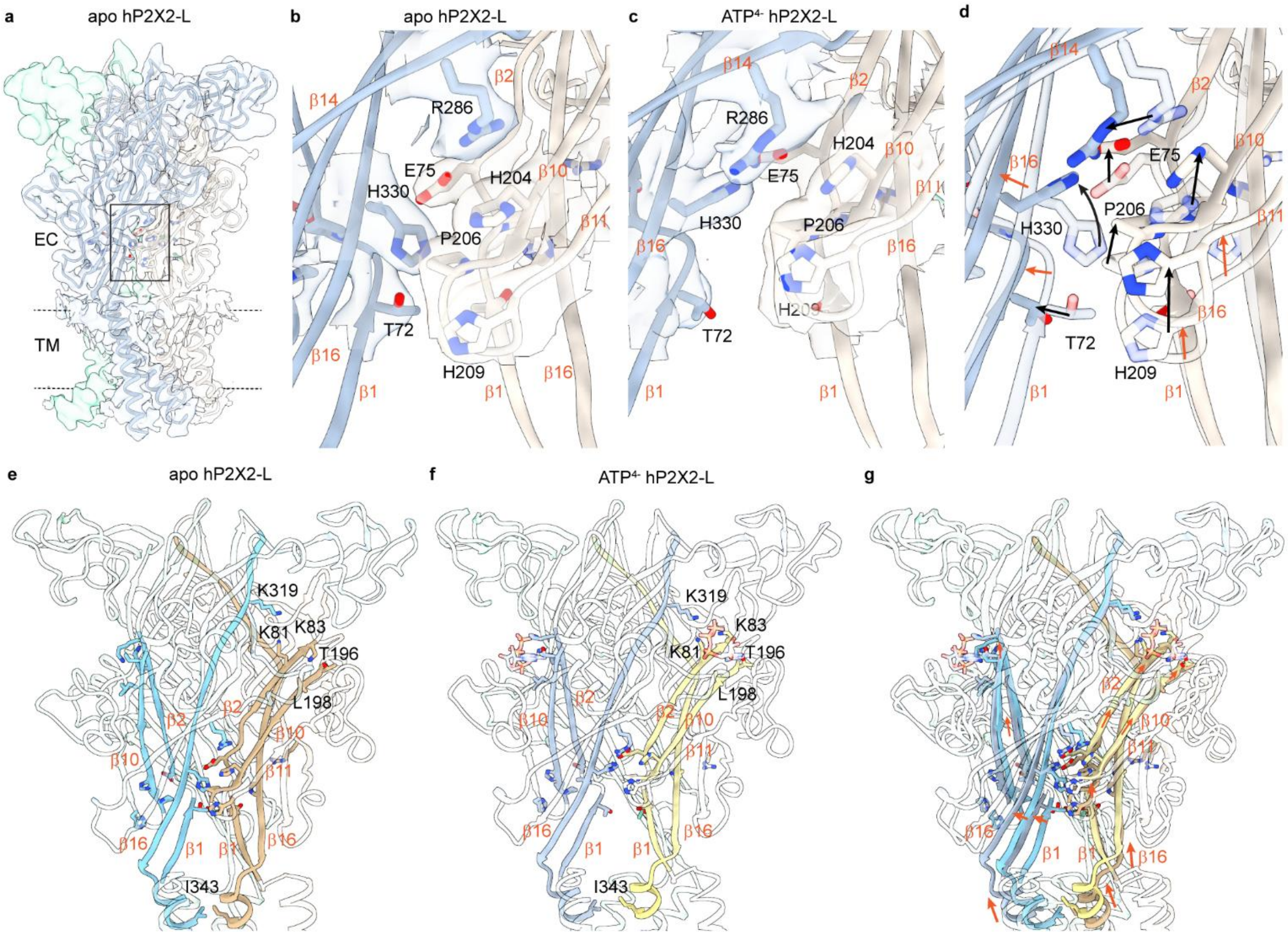


**Fig. S10. Structural elements involved in coupling ATP binding to channel opening.**
**a**) Cryo-EM map and structural model for apo hP2X2-L highlighting the location of a nexus of residues (boxed region) whose interactions change between apo, Mg-$ATP^{2-}$ and $ATP^{4-}$ bound. **b**) Cryo-EM density (DeepEMhancer sharpened) and structural model for apo hP2X2-L showing interactions between H330 and P206 and surrounding residues. **c**) Cryo-EM density (DeepEMhancer sharpened) and structural model for $ATP^{4-}$ bound hP2X2-L (class 2) showing interactions between H330 and E75. **d**) Superimposition of structural models for apo and $ATP^{4-}$ bound (class 2) hP2X2-L showing movement of residues between closed and open state. **e-g**) Overview of how the nexus connects to the ATP binding sites and the TM helices through a series of β-strands using the structure of apo hP2X2-L **(e)**, hP2X2-L with $ATP^{4-}$ bound (class 2, conformation 1) **(f)** and their superimposition **(g)**.

**Table S1. Cryo-EM data collection, refinement and validation statistics.**

| | hP2X2L-ND-apo (PDB- 9OGH, EMD-70468) | hP2X2L-DDM with residual ATP (PDB-9OIR, EMD- 70527) | hP2X2S-ND-Mg-ATP$^{2-}$ (PDB- 9Z32, EMD-73782) | hP2X2L-ND-Suramin (PDB-9OHK, EMD-70496) |
|---|---|---|---|---|
| **Data collection and processing** | | | | |
| Magnification | 105,000 x | | | |
| Voltage (kV) | 300 | | | |
| Electron exposure (e–/Å$^2$) | 51.42 | 58.22 | 59.34 | 60.80 |
| Defocus range (μm) | ~ -0.8 to ~ -2.4 | | | |
| Pixel size (Å) (Sup res.) | 0.415 | 0.411 | 0.412 | 0.412 |
| Symmetry imposed | C3 | | | |
| Initial particle images (no.) | 3,493,607 | 2,077,812 | 9,783,392 | 2,887,729 |
| Final particle images (no.) | 336,834 | 189,160 | 344,336 | 277,497 |
| Map resolution (Å) | 2.70 | 3.00 | 2.83 | 3.20 |
| FSC threshold | 0.143 | 0.143 | 0.143 | 0.143 |
| **Refinement** | | | | |
| Initial model used (PDB code) | AlphaFold 3 | 9ON5 | AlphaFold 3 | 9OGH |
| Model resolution (Å) | 2.7 | 3.0 | 2.9 | 3.2 |
| FSC threshold | 0.143 | 0.143 | 0.143 | 0.143 |
| Map sharpening *B* factor (Å$^2$) | -50 (Cryosparc) or DeepEMhancer (tightTarget / highRes models) | | | |
| Model composition | | | | |
| Non-hydrogen atoms | 8257 | 7869 | 7902 | 8127 |
| Protein residues | 990 | 978 | 972 | 987 |
| Ligands | NAG: 9<br>POV: 24<br>NA: 1 | NAG: 9<br>ATP: 3 | NAG: 9<br>ATP: 3 | NAG: 9<br>SVR:3 |
| *B* factors (Å$^2$) | | | | |
| Protein | 88.80 | 77.03 | 143.87 | 110.89 |
| Ligand | 121.54 | 50.81 | 126.67 | 141.87 |
| R.m.s. deviations | | | | |
| Bond lengths (Å) | 0.005 | 0.005 | 0.004 | 0.004 |
| Bond angles (°) | 1.064 | 1.078 | 0.928 | 1.131 |
| Validation | | | | |
| MolProbity score | 1.84 | 1.69 | 1.55 | 1.99 |
| Clashscore | 5.45 | 5.65 | 6.53 | 11.37 |
| Poor rotamers (%) | 2.12 | 0.48 | 1.56 | 0.35 |
| Ramachandran plot | | | | |
| Favored (%) | 95.73 | 94.44 | 98.14 | 93.58 |
| Allowed (%) | 4.27 | 5.56 | 1.55 | 6.42 |
| Disallowed (%) | 0 | 0 | 0 | 0 |

**Table S2. Cryo-EM data collection, refinement and validation statistics.**

| | hP2X2S-ND-apo (EMD-70457) | hP2X2S-ND-apo $Mg^{2+}$ (EMD- 70456) | hP2X2L-DDM with residual ATP after apyrase (EMD- 70526) |
|---|---|---|---|
| **Data collection and processing** | | | |
| Magnification | | 105,000 x | |
| Voltage (kV) | | 300 | |
| Electron exposure ($e^-/Å^2$) | 52.10 | 50.73 | 56.80 |
| Defocus range (µm) | | ~ -0.8 to ~ -2.4 | |
| Pixel size (Å) (Sup res.) | 0.415 | 0.412 | 0.411 |
| Symmetry imposed | | C3 | |
| Initial particle images (no.) | 1,828,228 | 1,820,541 | 2,285,521 |
| Final particle images (no.) | 190,930 | 234,490 | 162,474 |
| Map resolution (Å) | 2.93 | 2.97 | 2.80 |
| FSC threshold | 0.143 | 0.143 | 0.143 |

### Table S3. Cryo-EM data collection, refinement and validation statistics.

| | hP2X2L-ND-ATP$^{4-}$-Na$^{+}$ Class 1 State1/State2 (PDB- 9ON5/9ON6, EMD-70629) | hP2X2L-ND-ATP$^{4-}$-Na$^{+}$ Class 2 (PDB- 9OJK, EMD-70543) | hP2X2L-ND-ATP$^{4-}$-Na$^{+}$ Class 3 (PDB- 9OM0, EMD-70602) | hP2X2L-ND-ATP$^{4-}$-Na$^{+}$ Class 4 (PDB- 9OMR, EMD-70617) |
|---|---|---|---|---|
| **Data collection and processing** | | | | |
| Magnification | 105,000 x | | | |
| Voltage (kV) | 300 | | | |
| Electron exposure (e–/Å$^2$) | 59.90 | | | |
| Defocus range (μm) | ~ -0.8 to ~ -2.4 | | | |
| Pixel size (Å) (Sup res.) | 0.412 | | | |
| Symmetry imposed | C3 | | | |
| Initial particle images (no.) | 5,365,203 | | | |
| Final particle images (no.) | 106,536 | 172,339 | 243,758 | 65,876 |
| Map resolution (Å) | 2.60 | 2.60 | 2.58 | 2.70 |
| FSC threshold | 0.143 | 0.143 | 0.143 | 0.143 |
| **Refinement** | | | | |
| Initial model used (PDB code) | 9OJK | AlphaFold 3 | 9OJK | 9OJK |
| Model resolution (Å) | 2.6/2.6 | 2.6 | 2.6 | 2.7 |
| FSC threshold | 0.143 | 0.143 | 0.143 | 0.143 |
| Map sharpening *B* factor (Å$^2$) | N.A. DeepEMhancer (tightTarget / highRes models) | | | |
| Model composition | | | | |
| Non-hydrogen atoms | 7791 | 7743 | 7791 | 7584 |
| Protein residues | 969 | 963 | 969 | 945 |
| Ligands | NAG: 9<br>ATP: 3 | NAG: 9<br>ATP: 3 | NAG: 9<br>ATP: 3 | NAG: 9<br>ATP: 3 |
| *B* factors (Å$^2$) | | | | |
| Protein | 55.57/53.38 | 52.80 | 55.37 | 55.01 |
| Ligand | 73.53/70.91 | 70.18 | 71.59 | 70.23 |
| R.m.s. deviations | | | | |
| Bond lengths (Å) | 0.005/0.006 | 0.005 | 0.005 | 0.005 |
| Bond angles (°) | 1.017/1.010 | 0.987 | 0.996 | 0.981 |
| Validation | | | | |
| MolProbity score | 1.59/1.75 | 1.52 | 1.52 | 1.45 |
| Clashscore | 3.89/4.09 | 3.46 | 4.02 | 3.27 |
| Poor rotamers (%) | 1.81/2.53 | 1.82 | 2.17 | 1.49 |
| Ramachandran plot | | | | |
| Favored (%) | 96.57/96.26 | 96.87 | 97.61 | 96.81 |
| Allowed (%) | 3.43/3.73 | 3.13 | 2.39 | 3.19 |
| Disallowed (%) | 0 | 0 | 0 | 0 |

**Movie S1. Morph between apo and class 1 to 4 with $ATP^{4-}$ bound,** Morph between structures of apo hP2X2-L in the closed state and structures with $ATP^{4-}$ bound for class 1, class 2 (open), class 3 and class 4 (desensitized) constructed using ChimeraX. Key residues for binding ATP and those forming both activation and desensitization gates are shown in stick representation.

**Movie S2. Morph between apo and class 2 with $ATP^{4-}$ bound.** Morph between structures of apo hP2X2-L in the closed state and structures with $ATP^{4-}$ bound for class 2 (open) constructed using ChimeraX. Key residues involved in ATP binding along with those in β-strands involved in coupling $ATP^{4-}$ binding to channel opening are shown in stick representation.
